# Deconvoluting single-cell transcriptomics reveals cellular programs regulated by cell-cell communication in colorectal cancer

**DOI:** 10.1101/2025.04.09.648030

**Authors:** Han Zhang, Binfeng Lu, Aodong Qiu, Gregory F. Cooper, Anwaar Saeed, John W. Paisley, Xinghua Lu, Lujia Chen

## Abstract

Cells within a tissue microenvironment communicate through intricate cell-cell communication (CCC) networks. In this meta-analysis of eight single-cell cohorts encompassing 153 patients and 279 samples, we advance the understanding of CCC networks in colorectal cancers through a novel analytical framework. Employing hierarchical topic modeling, we identify gene expression modules (GEMs) that mirror single-cell signaling states, crucial for deciphering the complexity of intercellular interactions. By applying causal discovery methods, we systematically uncover GEMs likely regulated by ligand-receptor signaling and cross-cell-type communication. This analysis reveals cross-cell-type CCC programs, marked by highly correlated GEMs across various cell types, shedding light on the intricate CCC networks within the tumor microenvironment. Spatial transcriptomics further validate these findings by demonstrating the co-localization of GEMs within CCC programs in distinct spatial domains, emphasizing the spatial dynamics of tumor intercellular communication. Our interactive website (http://44.192.10.166:3838/) and analytical framework equip researchers with powerful tools to explore these complex mechanisms, potentially uncovering novel drug targets and refining strategies for precision immunotherapies. This comprehensive study not only presents a detailed catalog of CCC networks driven by ligand-receptor interactions in colorectal cancer but also highlights the significance of integrating multi-sample and patient data to unravel the molecular underpinnings of cancer communication pathways.

## Introduction

Colorectal cancer (CRC) is the 3^rd^ most common cancer and the 2^nd^ cause of cancer death ^1^. The discovery of the consensus molecular subtypes (CMS) among CRC tumors reflects the heterogeneity of composition and functional states of cells in the tumor microenvironments (TME), which in turn may underlie diverse behaviors of disease progression and responses to anticancer treatments by tumors. For example, except for those with deficient mismatch repair (dMMR), most CRCs do not respond to anti-PD-1 immune therapy ^2^. Such a relationship between genomic pattern and clinical response is the result of cell-cell communications (CCC) within the TME, through which the downstream signaling pathways are mediated and regulated ^3,4^. Therefore, a comprehensive understanding of the cell states and communication network in the TME are vital for uncovering the heterogeneous mechanisms of tumor growth, immune evasion and treatment resistance, which are essential for precision oncology.

Modeling CCC requires inferring the cellular states of individual cells, identifying synchronous changes in the cell-state among various cell types that may arise due to CCC, and investigating the molecular mechanism underlying CCC ^5–8^. Single-cell RNA sequencing (single-cell RNA-seq) is a powerful technology for profiling the transcriptomes of individual cells. As gene expressions are tightly regulated in a cell, the co-expression patterns of genes serve as proxies of the states of signaling pathways in a cell, which can be discovered by deconvoluting scRNA-seq data. Furthermore, the emergence of spatial transcriptomic technology provides additional spatial relationships between cells within their native spatial context ^9,10^. Thus, integrating scRNA-seq with spatial transcriptome provides the spatial context for interactions among different cell populations ^11^ and enhances the accuracy of modeling CCC.

Single-cell transcriptomes can be used to investigate cellular states at different levels. At the whole transcriptome level, cells sharing similar overall transcriptomic profiles can be discovered as discrete cell clusters using state-of-the-art clustering techniques ^12–15^. Although these approaches are good at classifying different cell populations, it does not allow examination of transcriptomic programs within individual cells that underpin the biological functions of these cells ^16^. To discover and represent transcriptomic programs within each cell, different statistical- and factorization- based models, e.g., non-negative matrix factorization (NMF) and latent Dirichlet allocation (LDA), have been used to deconvolute transcriptional programs by capturing the co-expression pattern of certain cell subsets ^17–20^. However, LDA models impose strong assumption of the independence among gene programs while NMF models do not explicitly identify dependencies among programs. Moreover, these methodologies fail to capture the well- established phenomenon that cells belonging to a cell type, such as T and NK (TNK) cells, exhibit highly coordinated transcriptomic programs. As a result, both approaches inadequately represent the nested nature of gene expression modules (GEM) that are observed in biology. These GEMs are more accurately represented as a hierarchical structure, wherein transcriptomic programs shared by cells across different cell subtypes, such as CD8+ T cells and CD4+ T cells, are positioned close to the root of the hierarchy. In contrast, transcriptomic programs underlying the differentiation of specialized cells are positioned as leaves. The methods of capturing such hierarchical relationships of transcriptomic programs remain relatively underdeveloped.

Transduction of signals across cells involves ligand-receptor interactions. Different methods have been developed to find potential ligand-receptor-mediated intercellular communication, e.g., CellChat, CellPhoneDBv2.0 ^6,21,22^. However, those methods are mainly based on the expression and enrichment of ligand-receptor pairs in specific cell clusters but do not explicitly model the impact of the signals transmitted by a ligand-receptor pair, e.g., genes regulated by PD-L1-PD-1 signaling. Furthermore, few methods exist for modeling coordinated cellular states of different cells within a tumor, e.g., certain subtype of myeloid cells and subtype of stromal cells tend to co-exist in tumors, or inferring whether the changed state of one type of cell directly influences the state of other types of cells.

To solve these challenges, we propose a novel framework for integrating bulk, single-cell, and spatial RNA-seq data to depict a comprehensive landscape of CRC TMEs (**Fig. 1**). We employed the nested Hierarchical Dirichlet process (nHDP) model to deconvolute a large collection of single-cell transcriptomes from 8 cohorts with over 153 CRC patients and over 626,858 single-cells (**Fig. 1A, Extended Data Table. 1 & Fig. 1**). nHDP organizes the discovered gene expression modules (GEMs) in a hierarchy, enabling it to separate GEMs (root node) representing general transcriptomic processes from those reflecting specific functional state of cells (leave node) (**Fig. 1B & 1C**). We then systematically investigated whether the expression of a GEM was regulated by ligand-receptor signaling (**Fig. 1D**) using the hill function that is used in pharmacology to measure the binding affinity between ligand and receptor. By studying the coordinated expression of GEMs, we systematically investigated the statistically significant intercellular communications across cell types (**Fig. 1E**) with spatial transcriptome and then extended our analysis to discover patterns of intercellular communication programs (GEM programs) shared within subpopulations of tumors (**Fig. 1F**). Employing techniques of causal analyses, we further searched for ligand-receptor pairs that contribute to mediate signal transduction of a CCC and further constructed a CCC network among cells expressing coordinated GEMs as well as biomarker discovery (**Fig. 1G**). We envision this comprehensive framework enables researchers to study complex mechanisms underlying distinct TMEs, investigate different immune evasion mechanisms, discover potential drug targets, and guide precision immune therapies.

**Fig. 1:**
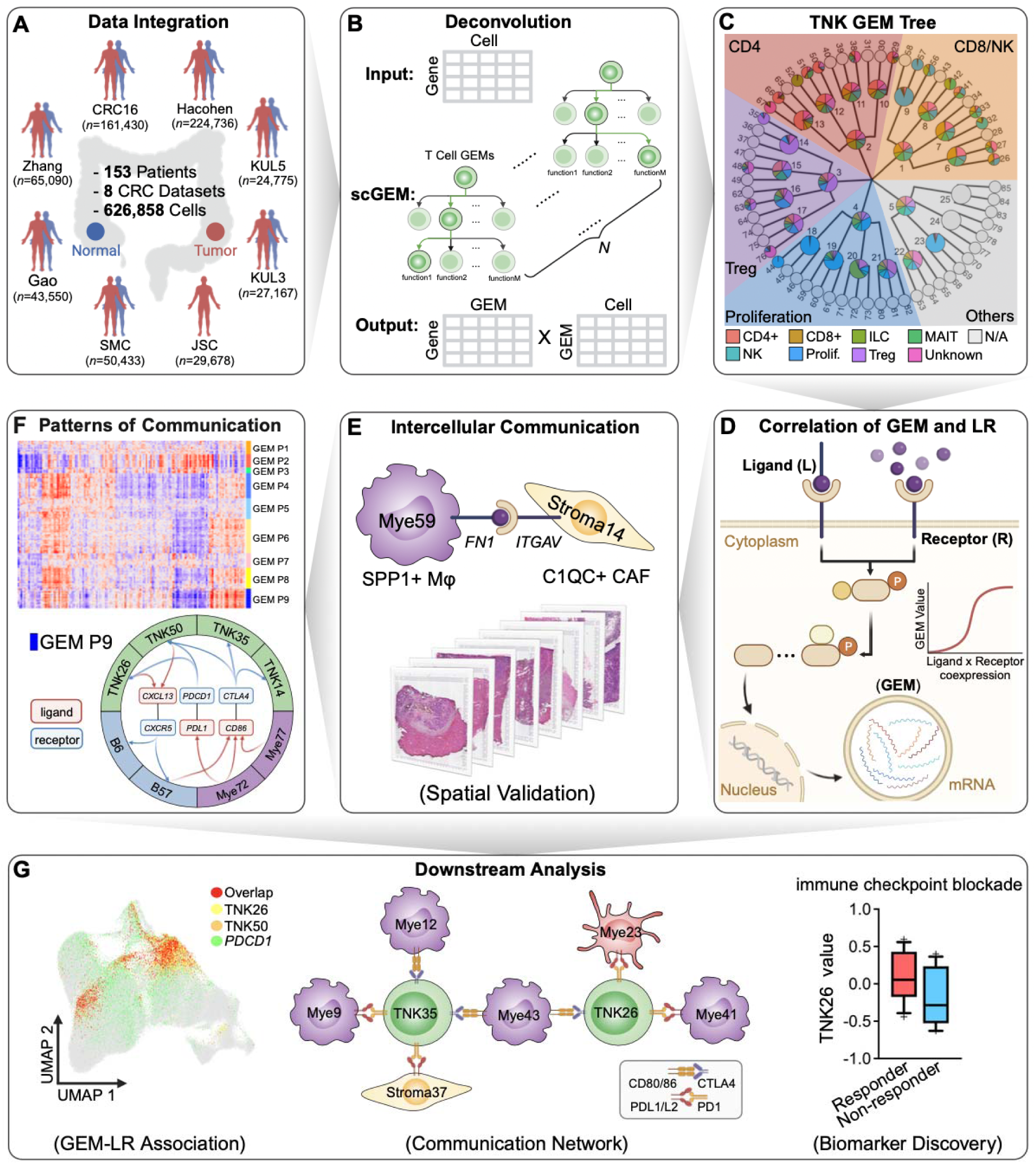
General design of the study. **A.** Merged CRC single-cell RNA-seq data from 8 cohorts. **B.** Diagram of the nHDP model using the expression of genes (a *cell-by-gene* matrix) in TNK cells as input. The transcriptome of one cell can be represented by a unique mixture of GEMs in the nHDP tree. The nHDP model outputs two matrices: a *gene-by-GEM* matrix reflecting the gene composition of a GEM and a *cell-by-GEM* matrix representing the GEM compositions of each cell. **C.** nHDP discovered the tree-structured organization of GEMs. Each node in the tree represents a GEM. A pie chart represents the percentage of each cell type the GEM is expressed. **D.** Illustration of modeling the relationship between ligand-receptor (LR) and its downstream GEMs. **E.** Modeling cell-cell communication by combining single-cell and spatial transcriptomes. **F.** Learning patterns of coordinated expression of GEMs across cell types to discover distinct GEM programs and associated tumor microenvironments. **G.** Downstream discoveries enabled by deconvoluting single-cell transcriptomes by nHDP: **1)** GEMs highly associated with the signal of LR. **2)** Subnetwork of intercellular communication for GEMs and LRs of interest. **3)** GEM as potential novel biomarker for therapies.

### Result

### A comprehensive atlas of GEMs expressed by cells in CRC tumor microenvironments

We collected single-cell RNA-seq data from 8 independent CRC cohorts. Then, we separated cells into 5 major cell types: TNK: mainly including T and natural killing (NK) cells (N=246,560); Myeloid: myeloid cells, including monocytes, macrophages, neutrophils, and dendritic cells (N=94,663); PlasmaB: including B and plasma cells (N=96,637); Stroma: stromal cells, including endothelial, fibroblast and myofibroblast cells (N=69,128); Epi: epithelial cells (N=119,870) (**Extended Data Table. 1**). The merged cells for each cell category were clustered and labeled using Seurat pipeline and canonical marker genes. After the integration and batch removal, we identified 18 clusters in TNK cells (**Fig. 2A**), 18 cell populations in both Myeloid cells and PlasmaB cells and 17 cell groups in Stroma cells (**Extended Data Fig. 1**). We also showed that the distributions of datasets are well maintained in all clusters in TNK cells (**Fig. 2B**), suggesting the batch effects were regressed out; thus, the pooled TNK cells reflect the landscape of tumor-infiltrating TNK cells in CRC.

**Fig. 2:**
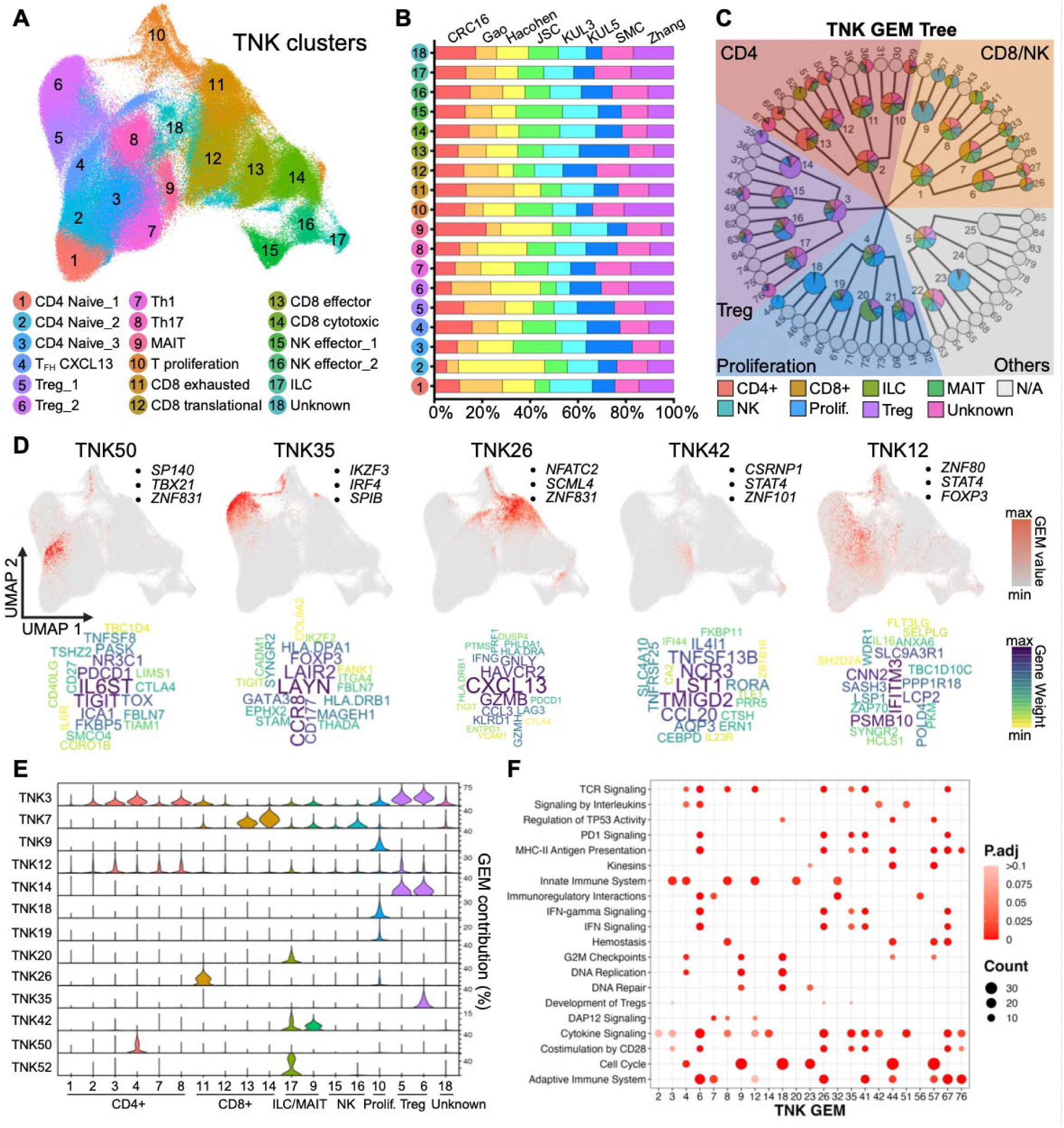
Deconvoluting single-cell transcriptomics to discover GEMs reflecting the state of signaling pathways. **A.** UMAP plot of 246,560 merged TNK cells from 8 CRC cohorts. **B.** The proportion plot reflects the origins of cells in each cell cluster. **C.** The hierarchical tree structure of TNK GEMs was learned from the nHDP model, which was trained with merged TNK cells. The nHDP model has three levels. The first level contains GEMs from 1-5. The second level contains GEMs from 6-25. The third level contains GEMs from 26-85. Each color represents a subtype of T cell. The unused GEMs are colored grey. **D.** The UMAP plot, word cloud plot, and top associated TFs of selected TNK GEMs. **E.** The functional involvement of TNK GEMs within each cell cluster. **F.** The enriched signaling pathways associated with TNK GEMs.

To model the tree structure of the TNK gene expression modules (GEMs), we utilized the nested hierarchical Dirichlet process (nHDP) model to identify a series of nested gene expression modules (GEMs) ^23^, where a GEM is a probability distribution over the gene space that captures the coordinated expression pattern of a set of genes in a subpopulation of cells. The nHDP model organizes inferred GEMs in a hierarchical tree structure, and we set branching factors of 5, 4, and 3 at 1^st^, 2^nd^, and 3^rd^ layers, respectively, leading to a total of 85 nodes to capture GEMs with different levels of granularity and specificity. **Fig. 2C** shows the tree organization of the GEMs learned using TNK cells as input. The nHDP model populated the nodes in the hierarchy through a data-driven approach. Each node in the tree represents a GEM. The ones close to the root (TNK GEMs 1-5 in **Fig. 2C**) represent GEMs that are more broadly expressed in all T, innate lymphoid cells (ILC), mucosal-associated invariant T cells (MAIT) and NK cells, which can be used to define major subtypes of T and NK (TNK) cells. The ones close to the leaves represent GEMs specifically expressed in more differentiated cells. In the TNK GEM tree, only 43 TNK GEMs were identified as biologically meaningful and represented as filled nodes. The nodes identified as biologically meaningless were shown as empty nodes without color.

We systematically inspected the TNK GEMs reflecting TNK transcriptomic processes. The pie chart within each GEM illustrates the proportion of cells expressing that specific GEM across various T cell subtypes, including CD8^+^ T cells, CD4^+^ T cells, ILC, MAIT, NK cells, regulatory T cells (Treg) and Proliferation T cells. It is interesting to note that major branches of the nHDP hierarchy tree, as defined by GEMs 1-4, represent major TNK subtypes, including CD8/NK, CD4, Treg, and proliferation T cells (**Fig. 2C**). Projecting the expression of GEMs onto the UMAP of TNK cells (**Fig. 2D**), we noticed that certain GEMs were concentrated in a particular cell cluster or a combination of clusters, representing a characteristic transcriptomic program that is active in the cell cluster. For example, TNK50 is concentrated in cells belonging to cluster #4, which consists of T helper (Th) cells expressing CXCL13; TNK35 is mainly expressed in T cell cluster #6, a subset of T_reg_. Conversely, there are also GEMs shared across multiple cell clusters, suggesting a transcriptomic program that is shared across multiple subtypes of T cells. For example, TNK26 is expressed in a subset of CXCL13^+^CD4^+^ T cells (cluster #4), MKI67^+^ proliferative T cells (cluster #10), and CXCL13^+^CD8^+^ T cells (cluster #11). TNK42 is expressed in MAIT cells (cluster #9) and ILC (cluster #17). Finally, TNK12 is expressed broadly among Treg, Th, and proliferating T cells. These results indicate that our model can capture transcriptomic programs that are both shared and specialized by cells belonging to different cell subtypes. To test the performance of our model, we also compared the distributions of the transcriptomic programs that were found by our model and NMF ^24^. Our result demonstrates that the nHDP excels in capturing more specific cellular programs shared across multiple cell clusters when compared to NMF as shown in **Extended Data Fig. 2**. For instance, TNK15 captures a cellular program of greater specificity, shared among cells from 4 TNK cell clusters, surpassing the performance of NMF5. NMF5 captures a more generalized cellular program that combines programs represented by TNK15, TNK42, and TNK51. Similarly, NMF58 captures the cellular programs combining programs represented by TNK17, TNK29, and TNK50.

To make the GEMs more biologically explainable, we depict the distinctive gene composition profile of each GEM (top 20) through a word cloud, and the top three associated transcription factors identified by transcription factor analysis using ChEA3 ^25^ were illustrated as black dots in **Fig. 2D**. It is interesting to note that TNK26 is enriched with genes characteristic of exhausted T cells including PDCD1, CXCL13 and TIGIT. Based on the transcription factor analysis, NFATC2 regulated the top 50 genes associated with TNK26. Several studies have shown that the transcription factor NFAT is involved in the exhaustion process of activated CD8^+^ T cells ^26^. In addition, the top genes associated with TNK26 and TNK50 are both regulated by transcription factor ZNF831, which has been shown to inhibit the functional activity of effector cells through STAT pathway ^27^. This indicates the potential existence of a signal leading to the regulation of T cell exhaustion. We further examined the distribution of GEMs in different TNK clusters (**Fig. 2E**), unveiling which GEMs are broadly shared among cells assigned to multiple clusters and which GEMs are specific to a particular cell cluster. **Extended Data Fig. 3** displays the detailed comparison between GEMs and TNK cell clusters by measuring the correlation between top 50 genes of a GEM and highly variable genes in each TNK cell subtype. Furthermore, we performed pathway enrichment analyses in Reactome to investigate the functions of top genes associated with GEMs (**Fig. 2F**). Interestingly, several TNK GEMs, including TNK6, TNK26, TNK35, and TNK41, are associated with PD-1 signaling pathway. The dot plots showing the distribution of GEMs in traditional cell clusters and the enriched signaling pathways for other cell types (PlasmaB, Myeloid, Stromal) are listed in **Extended Data Fig. 4 & 5**. The detailed annotation of each GEM, including the UMAP, cell composition, and top associated genes, is listed in our interactive website. Besides, the enriched MsigDB gene sets (H & C1-C8) for each GEM, derived from the hypergeometric analysis, is shown in **Extended Data Table. 7**.

In summary, by deconvoluting the transcriptomes of a large number of single cells using nHDP, our approach revealed GEMs that represent transcriptomic programs within or across traditional cell subtypes, which reflect certain signaling events within these cells.

### Discovering potential regulatory relationships between ligand-receptor signaling and GEMs

We set out to examine if the expression of a GEM in cells is potentially regulated by certain ligand-receptor signaling by testing the hypothesis posited in **Fig. 3A**: If a ligand-receptor pair regulates the expression of the member genes of a GEM, the overall expression value of the GEM can be modeled with a sigmoid dose-response model with respect to ligand-receptor pair. We modeled the relationship of the expression of ligand, receptor, and GEM at the tumor sample level. We first generated pseudo-bulk RNA-seq (a *tumor-by-GEM* matrix) from the single-cell RNA-seq (a *cell-by-GEM* matrix) for tumor samples. Then we estimated the concentration of ligand and receptor proteins based on the pseudo-bulk expression (see Methods) of the ligand ([*L*]) and receptor ([*R*]) for each of the 151 tumors. Additionally, we approximated the concentrate of protein complex as the product of ligand and receptor: [*LR*] ∝ [*L*]*[*R*] ^5^. Pseudo- bulk expression of GEMs was estimated using the GSVA analysis. Subsequently, we conducted a systematic search against all GEMs with known LR pairs from NicheNet database and retained the LR pairs that could predict the expression of the GEM with statistical significance (**Extended Data Table. 2**).

**Fig. 3:**
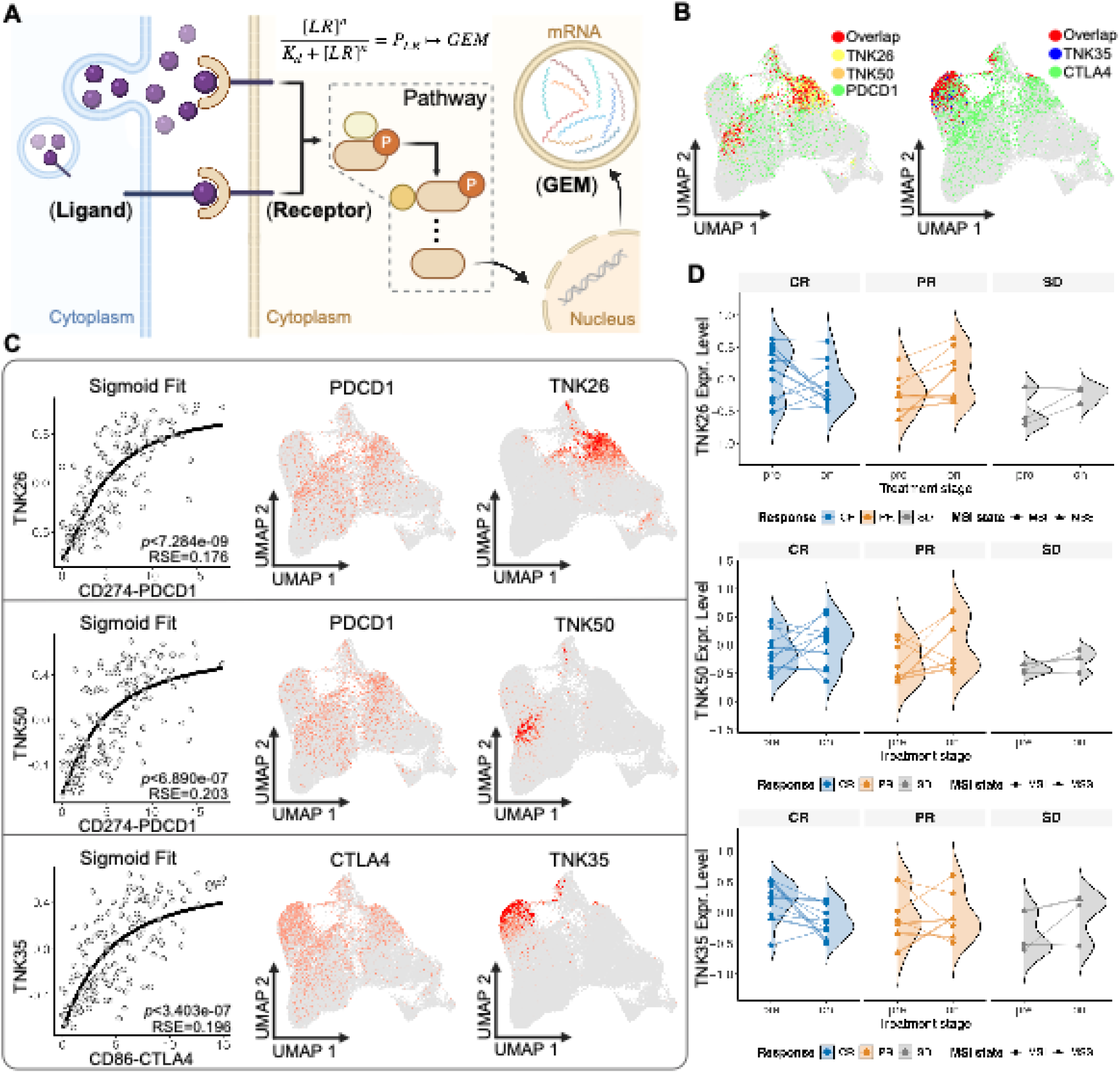
Discovering ligand-receptors strongly associated with specific pathways regulating the expression of GEMs. **A.** Illustration of GEMs reflective of downstream transcriptome mediated by biochemical signaling through ligand-receptor binding. The significant association between a GEM and a ligand-receptor pair needs to satisfy the sigmoid fitting. **B.** The overlap of cells expressing receptor and GEMs of interest. Left: the overlap of *PDCD1*, TNK26 and TNK50. Right: the overlap of *CTLA4* and TNK35. **C.** From left to right, the plot of sigmoid fitting between GEM and ligand-receptor pair with *p*-value and residual standard error (RSE), UMAP plot of the receptor gene, and UMAP plot of the GEM. **D.** Comparison of the expression levels of TNK26, TNK50 and TNK35 in the CR (complete response), PR (partial response) and SD (stable disease) groups before (pre-) and during (on-) anti-PD-1 immunotherapy treatment.

We investigated the GEMs that are strongly associated with immune checkpoints programmed cell death protein 1 (PD-1, encoded by *PDCD1*) and cytotoxic T lymphocyte-associated protein 4 (*CTLA4*) signaling (**Fig. 3B**). For the PD-1 signaling, the top two TNK GEMs that could be predicted by approximated [*LR*] with the strongest non-linear correlation were TNK26 and TNK50, with Spearman’s correlation coefficients between prediction and observation being *ρ* =0.76, *p*<2.2e-16 and *ρ* =0.68, *p*<2.2e-16, respectively. The UMAPs show that the distributions of *PDCD1* and the two GEMs are highly correlated at the single-cell level (**Fig. 3B-C**). The results also suggest potential PD-L1-PD-1 signaling is associated with distinct GEMs in different T cell subtypes. Specifically, [*CD274*]*\**[*PDCD1*] is associated with TNK50 in CXCL13^+^CD4^+^ T cells, whereas [*CD274*]*\**[*PDCD1*] is associated TNK26 mainly in exhausted CD8^+^ T cells. Importantly, the approximated ligand-receptor complex [*LR*] is more predictive of the expression of these two GEMs when compared with only using the receptor expression [R] alone. Similarly, the results show that the expression values of TNK35 can be predicted by approximated *CD86* and *CTLA4* complex with Spearman’s *ρ* =0.59, *p*<2.2e-16. The expression of *CTLA4* is also highly overlapped with the expression of TNK35 in Treg cells (**Fig. 3B-C**), suggesting the formation of this complex at these cells is associated with expression of the GEM in a non-linear dose-response fashion. In summary, our results show that identifying GEMs enabled the investigation of the functional impact of potential ligand-receptor signaling. We visualized a subset of ligand-receptor pairs and TNK GEMs that exhibited statistically significant nonlinear dose-response relationships (**Extended Data Fig. 6 & Table. 2**).

The strong non-linear dose-response relationship between a ligand-receptor pair and a GEM (e.g., *CD274-PDCD1* and TNK26) suggests the hypothesis that the expression of some members of a GEM may be regulated by the ligand-receptor interaction. Thus, PD-1 signaling pathway is likely to be more active in tumors expressing a high level of TNK26, and thereby, these tumors might be more susceptive to anti-PD-1 treatment. To test the association of the expression level of TNK26 and response to anti-PD-1 related treatment, we collected clinical and transcriptomic data from Chen et al.’s CRC cohort treated with anti-PD-1 plus chemotherapy ^4^, which involves 20 CRC patients. The transcriptome and clinical responses to the treatment were measured pre-, on-, and post-treatment. The patients were classified as CR (complete response), PR (partial response), and SD (stable disease), based on the classification provided in the study. We compared the expression level of GEMs learned from nHDP model in these three groups using the pre-treatment transcriptome to identify discriminative GEM features. The analysis indicates a higher TNK26 expression associated with CR group compared to the PR and SD group (**Fig. 3D**). This association was further tested across one established CRC studies involving patients treated with the combination of carbozantinib (a small molecule inhibitor of tyrosine kinases including VEGFR2) and durvalumab (an immune checkpoint inhibitor targeting PD-L1) ^28^ and three established melanoma studies ^29–31^ involving patients treated with anti-PD-1 antibodies. We performed the wilcoxon test to identify discriminative GEMs. Consistently, TNK26 expression is significantly higher in the responder group compared to the non-responder group (**Extended Data Fig. 7A & 7C**). Moreover, by comparing the peak of TNK26 expression distribution before and after treatment (**Fig. 3D**), we observed an apparent decrease following the anti-PD-1 therapy, suggesting that TNK26 may be a downstream target of PD-1 signaling in the CR group. This discovery is aligned with the statement from the original paper that CR patients experience a transition from a hot to a cold immune state driven by effective tumor clearance. The results support the notion that the expression level of TNK26 reflects the involvement of the PD-1 signaling in tumors, and thus can potentially be used as a novel biomarker for responsiveness to the anti-PD-1 therapy. Regarding TNK50, a significantly higher expression in responder group than in non-responder group is observed exclusively in the melanoma dataset. In the CAMILLA trial, even though TNK50 is shown to be correlated with TNK26, TNK50 did not reach statistical significance, likely due to the small number of responders with one apparent outlier, as depicted in **Extended Data Fig. 7B**. More examples of GEMs showing significant difference between responder group and non-responder group in CAMILLA trial and melanoma cohorts are listed in **Extended Data Fig. 7A & 7C**, respectively.

### The CCC between SPP1-macrophage and cancer-associated fibroblast through ligand-receptor *SPP1-ITGAV*

In addition to the cell-cell communications of TNK cells in the TME, the significance of crosstalk between macrophages and cancer-associated fibroblasts (CAFs) has garnered increasing attention in recent studies ^32–34^. Motivated by this, we embarked on an investigation into this crucial aspect. After searching for highly correlated GEMs that are expressed in stromal cells and myeloid cells, our analysis led to a pair of GEMs, Mye11-Stroma11, which exhibited strong Spearman correlation (ρ = 0.35, p = 9.2e-06) at the pseudo-bulk RNA level (**Fig. 4A**). The results led to a hypothesis that myeloid cells expressing Mye11 may communicate with CAF cells expressing Stroma11. Based on the annotation of GEM distribution, Mye11 is expressed in a separate subpopulation of M2 macrophage, Stroma11 is expressed in a subpopulation of tumor promoting CAF. After searching for the *LR* potentially mediating such communication, we identified a LR pair *SPP1/ITGAV*, which predict the expression of Stroma11 (**Fig. 4A**) in a nonlinear dose-response fashion (p-value of sigmoid fitting being 0.0004). Examining the distribution of the ligands and receptor genes and the above GEMs in UMAPs showed that *SPP1* and Mye11 were co-expressed in a subpopulation of M2 macrophage subpopulation (**Fig. 4B**). *ITGAV, which* encodes subunits of integrins, is widely expressed in stromal cells, including both fibroblasts and endothelial cells. This suggests that integrin signaling may have distinct downstream effects in different stromal subpopulations, with Stroma11 being one of them. This claim is supported by the existence of other *GEM X-LR-GEM Y* triplet communications guided through *ITGAV* (**Extended Data Table. 6**).

**Fig. 4:**
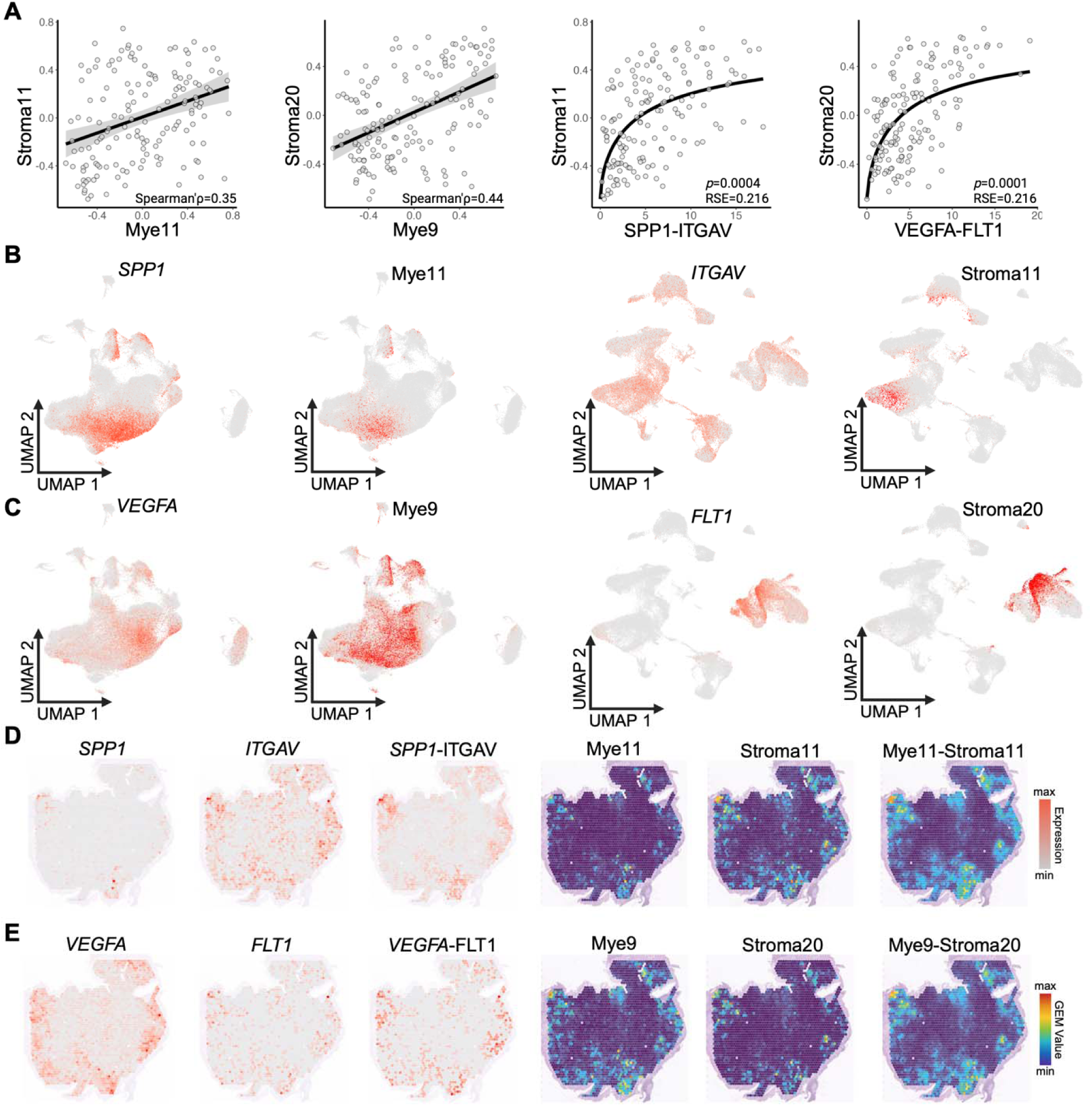
Modeling intercellular communication. **A.** Left two: The scatter plot of Mye11-vs-Stroma11 and Mye9-vs-Stroma20 using the pseudo-bulk transcriptome data; Right two: The sigmoid fitting between Stroma11 and ligand-receptor pair *SPP1-ITGAV*, and the sigmoid fitting between Stroma20 and ligand-receptor pair *VEGFA-FLT1* using the pseudo-bulk transcriptome data. B. The UMAP plot of cells expressing *SPP1*, Mye11, *ITGAV* and Stroma11. **C.** The UMAP plot of cells expressing *VEGFA*, Mye9, *FLT1*, and Stroma20. **D**. The spatial distribution plot of *SPP1*, *ITGAV*, *SPP1-ITGAV*, Mye11, Stroma11, and Mye11-Stroma11 using the spatial slide (SN048_A121573_Rep2) as an example. **E.** The spatial distribution plot of *VEGFA*, *FLT1*, *VEFGA-FLT1*, Mye9, Stroma20, and Mye9-Stroma20 using the same slide.

To reduce the false positive discoveries, we require the *GEM X-LR-GEM Y* triplet to pass both conditional independence test and spatial colocalization test. Based on the conditional independence test, Mye11 and Stroma11 become independent when conditioned on the presence of *SPP1-ITGAV*, indicating a potential causal relationship. We further tested this triplet using two spatial transcriptomic cohorts (metastatic CRC patients from Wu et al ^35^ and CRC patients from Valdeolivas et al ^36^) by analyzing the co-localization of the ligands, receptors, GEMs and their interactions in the spatial domains. The HE slides in these two spatial cohorts are provided in **Extended Data Fig. 10**. Using a CRC tumor section (SN048_A121573_Rep2) as an example, the spatial expression of ligand (*SPP1*), receptors (*ITGAV*) and GEMs (Mye11 and Stroma11) were highly correlated (**Fig. 4D**), indicating that cells expressing these genes and GEMs likely interact within spatial domains. The spatial correlation of Mye11-Stroma11, Mye11-*SPP1*, Stroma11-*ITGAV*, and Stroma11-*SPP1-ITGAV* were 0.78, 0.75, 0.41, and 0.51, respectively. The p-values associated with the spatial correlations shown above are less than 1e-22 (See **Extended Data Table. 6**). The spatial colocalization of this communication (Mye11-*SPP1/ITGAV*-Stroma11) was observed across multiple spatial slides (14 slides). These results support the hypothesis that SPP1+ macrophages communicate with CAFs via the *SPP1-ITGAV* ligand-receptor pair.

### The CCC between macrophage and endothelial cells through ligand-receptor *VEGFA-FLT1*

By analyzing the transcriptomic data from the CAMILLA trial involving combinational anti-PD-L1 immunotherapy, Stroma20 emerged as the most discriminative GEM feature, showing strong potential for predicting drug response in metastatic CRC patients (**Extended Data Fig. 7A**). Stroma20 is expressed in a subpopulation of tip cell/angiogenic endothelial cells. The top three associated TFs are BCL6B, SOX18, and SOX7 and the top enriched signaling pathway associated with the leading genes of Stroma20 is NOTCH1 signaling. The relationship between NOTCH signaling and VEGFR has been demonstrated in previous studies ^37^.

We then searched for the triplet pairs including Stroma20 that passed both the conditional independence test and spatial colocalization test, and identified one significant communication: Mye9-*VEGFA-FLT1*-Stroma20. Mye9 is a subpopulation of macrophage and dendritic cells, and exhibited strong correlation with Stroma20 (Spearman ρ = 0.44, p = 2.7e-08) at the pseudo-bulk RNA level (**Fig. 4A**). UMAPs in **Fig. 4C** show the co-expression of *VEGFA* and Mye9, and the co-expression of *FLT1* and Stroma20. The spatial expression of ligand (*VEGFA*), receptors (*FLT1*) and GEMs (Mye9 and Stroma20) were highly correlated, as shown in **Fig. 4E**. Using the same slide in **Fig. 4D**, the spatial correlations for Mye9-Stroma20, Mye9-*VEGFA*, Stroma20-*FLT1*, Stroma20-*VEGFA-FLT1* are 0.69, 0.56, 0.31, and 0.31, respectively. The p-values associated with the spatial correlations shown above are less than 1e-22 (See **Extended Data Table. 6**). This communication (Mye9-*VEGFA-FLT1*-Stroma20) is spatially colocalized in multiple slides (six slides). Our findings provide insights into identifying potential biomarkers and their associated communications for current and novel treatments that elucidate underlying biological mechanisms.

The example discussed above demonstrates our ability to identify potential CCC candidates mediated by the ligand-receptor signaling, supported by the integration of scRNA-seq (co-expression), pseudo-bulk RNA-seq (correlation and conditional independence test) and spatial transcriptomic data (spatial colocalization test).

### Discovering coordinated CCC programs

We further investigated patterns of coordinated expression of GEMs across different cell types (TNK, B, myeloid, stromal, and epithelial) in CRCs. We represented each tumor in the GEM space (pseudo-bulk expression: *cell-by-GEM* matrix) and performed double clustering of GEMs and tumors. The results (**Fig. 5A**) showed that the expression of many GEMs was highly coordinated. We hypothesized that the coordinated expression of GEMs across multiple types of cells in a TME is likely due to two possible mechanisms: environmental factors that orchestrate the expression of GEMs (confounding) or a CCC network that allows cells to communicate and reach homeostasis. We refer to a set of GEMs across multiple cell categories that displays a highly coordinated expression pattern as a CCC program, and we have identified 9 CCC programs (rows). Based on their GEM composition, clustering tumors revealed 6 sample clusters (columns). A complete list of CCC programs is in **Extended Data Table. 4**.

**Fig. 5:**
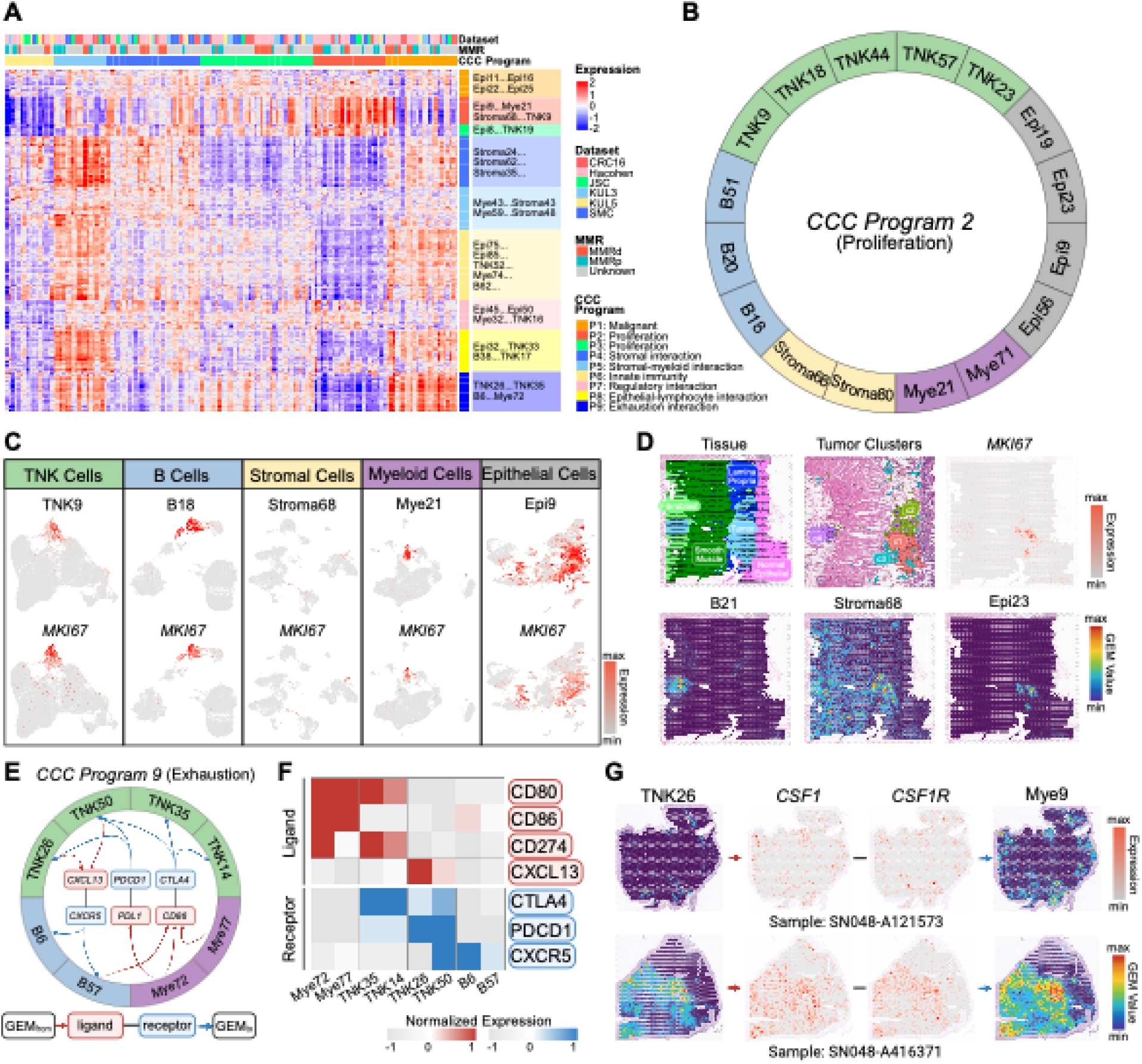
Discovering patterns of intercellular communication channels that define transcriptome programs across different cell types. **A.** Clustering of GEMs and CRC tumors. Pseudo-bulk profile for each tumor was constructed from single-cell RNA-seq data. Expression values of a GEM were estimated based on the GSVA using the top 50 weighted genes in a GEM. The GEMs (rows) are grouped into 9 CCC programs by clustering analysis, representing patterns of intercellular communication channels. The tumors (columns) are grouped into 6 clusters. **B.** The GEM members of the CCC Program 2. The majority of genes associated with these GEMs are MYC pathway targets, as indicated by the HALLMARK_MYC_TARGETS in GSEA. **C.** UMAP of MKI67 and GEMs in CCC program 2 in different cell-type categories. **D.** Spatial colocalization of proliferation marker MKI67 and GEMs in CCC Program 2. **E.** The potential CCC network and the ligand-receptors involved in GEMs in CCC Program 9. The red and blue rectangle in the center represents ligands and receptors respectively. The edge between a ligand/receptor and a GEM is added if the ligand/receptor is expressed in the cells expressing a GEM. **F.** The expression level of receptors (blue) and its associated ligands (red) in cells expressing the GEMs in CCC program 9. TNK and B GEMs in CCC Program 9 mainly express receptors (*PDCD1, CTLA4, and CXCR5*). Myeloid GEMs mainly express ligands (*CD80, CD86 and CD274*). **G.** The spatial colocalization of ligand, receptor and GEMs: TNK26 -> *CSF1-CSF1R*-> Mye8 in two spatial slides (SN048_A416371_Rep2 and SN048_A121573_Rep2).

We further investigate the potential molecular basis of CCC programs. CCC Program 2 consists of 16 GEMs from different cell categories (**Fig. 5B**). The pathway analysis indicates that GEMs in CCC Program 2 are significantly enriched of genes involved in cell cycle pathways, such as the C-MYC signaling pathway, a hallmark pathway of cell proliferation in different cell types ^38^. The GEMs belonging to this CCC program include GEMs from TNK, B, myeloid, epithelial and stromal cells, respectively (**Fig. 5B**), suggesting they are subpopulations of cells in a proliferation stage from each cell category. Of interest, we noted that TNK9 is enriched in the T- cell cluster #10, which is labeled as proliferative T cells. Not surprisingly, an epithelial GEM belonging to this program, Epi9, is broadly active in epithelial cells, reflecting a shared transcriptomic program involved in cell proliferation by cancer cells. The UMAPs in **Fig. 5C** indicate that various cell categories expressing these GEMs also express the proliferation marker gene *MKI67*. We hypothesized that the expression of GEMs of proliferation nature in different cell types may be associated with signals that promote proliferation of cells, e.g., receptor tyrosine kinase (RTK) pathways, Wnt/β-catenin pathway, and PI3K-AKT-mTOR pathway. Our pathway enrichment analysis shows that the top genes associated with most of GEMs in CCC Program 2 are enriched in the MYC pathway, which is downstream of RTKs. To test the hypothesis, we investigated the spatial distribution of the GEMs in tumors. The results (**Fig. 5D**) show the co-localization of GEMs in the CCC Program 2. We calculated the spatial correlations between MKI67 with respect to B21, Stroma68, and Epi23, using their expression values in spatial spots. The correlation coefficients are 0.708, 0.762, and 0.804, respectively, providing evidence for the strong coordinated expression of these GEMs in spatial domains. We further checked the cell composition of spatial spots expressing these GEMs. The inference of cell composition in tumor spots (**Extended Data Fig. 8A**) shows that those spots contain a mixture of cells potentially expressing these GEMs. The compositions of B21, Stroma68 and Epi23, represented by their top associated genes, are not overlapping (**Extended Data Fig. 8B**). Each of the three GEMs captures cell-type-specific proliferation signals. The indicates that the diverse cells expressing GEMs in GEM program 2 may be stimulated by proliferation-related factors existing in tissue microenvironments.

Moreover, we performed deconvolution analysis on the GEMs in CCC Program 2 utilizing the single-cell RNA-seq transcriptome data obtained from the peripheral blood sample prior to the treatment of the regorafenib and nivolumab combination therapy ^39^. Our analysis revealed that the expression levels of GEMs, including TNK9, B18, and Mye21, in the proliferation program (CCC program 2), exhibited significant differences between responders and non-responders, as illustrated in **Extended Data Fig. 9**. This observation aligns with the findings reported in the original paper and highlights the potential of cellular signals related to MKI67 proliferation as indicative markers for predicting clinical outcomes in patients treated with a combination of multi-kinase inhibitors and immune checkpoint inhibitors.

We also searched for CCC programs in which coordinated expression of GEMs is likely due to CCC between the GEMs. As an example, CCC Program 9 includes TNK26, TNK35, TNK50, Mye72, Mye77, B57, and B6. To investigate whether cells expressing these GEMs communicate with each other, we searched for LR pairs that were co-expressed with these GEMs, e.g., (*GEM X* and *L*) and (*R* and *GEM Y*), as a candidate pair of communicating cells, using the pseudo-bulk RNA-seq data. We found three main LR pairs through which GEMs in the program may communicate, including *CD274-PDCD1*, *CD86-CTLA4*, and *CXCL13-CXCR5*, which are closely associated with T-cell immune checkpoint molecules in the tumor microenvironment. We illustrate the possible communication channels among the cells expressing those GEMs using directed edges based on the co-expression of *GEM X-L* and *R-GEM Y* (**Fig. 5E**). The expression level of the ligands and receptors in the cells expressing GEMs in the CCC Program 9 is shown in **Fig. 5F**. The cells expressing TNK GMEs in CCC Program 9, such as TNK26 and TNK35, mainly express receptor genes *PDCD1* and *CTLA4*. GEM B57 is concentrated in IgA/G cells expressing the *CXCR5*, which is the receptor of *CXCL13*. Two myeloid GEMs, including Mye72 and Mye77, concentrated in dendritic cells, express ligand genes *CD274*, *CD80*, and *CD86*.

We then investigated how GEMs in CCC program 9, such as TNK26, communicate with GEMs in other CCC programs. To reduce false-positive discoveries, we required the communication triplets (*GEM X–LR–GEM Y*) to pass both the conditional independence test and spatial colocalization. **Fig. 5G** highlights one of the most significant communications involving TNK26 that meets both criteria: TNK26—*CSF1-CSF1R*—Mye8. The Spearman correlations of TNK26- Mye8, TNK26-*CSF1*, *CSF1R*-Mye8 in pseudo-bulk are 0.33 (*p*=3.5e-05), 0.39 (p=8.6e-07) and 0.57 (p<2.2e-16). The ligand, receptor, and two GEMs were shown to be colocalized in spatial domains in 11 slides. **Fig. 5G** uses two slides as examples to show the significant colocalization. The spatial correlations of TNK26-Mye8, TNK26-*CSF1*, and *CSF1-CSF1R*-Mye8 are 0.85, 0.78, and 0.60 (upper slide: SN048_A416371_Rep2), and 0.64, 0.32 and 0.39 (lower slide: SN048_A121573_Rep2), respectively. The p-values associated with the spatial correlations above are all less than 1e-22 (See **Extended Data Table. 6**). Several studies have demonstrated the importance of CSF1R signaling in CD8 T cell exhaustion in cancer types such as breast cancer ^40–42^, as well as the potential of CSF1R inhibitors as monotherapy or in combination with other immunotherapies, including immune checkpoint inhibitors ^43^. The strong association of TNK26 with signaling pathways, such as *PD-L1-PDCD1* and *CSF1-CSF1R,* highlights the potential of TNK26 and its related interactions as targets for mono/combination immunotherapy.

In summary, the findings presented in this section collectively illustrate the feasibility of uncovering a unique CCC mechanism within a micro-TME. This is achieved by modeling the synchronized expression of GEMs, identifying potential LRs responsible for mediating communication, and providing evidence of spatial colocalization for the components involved in the identified CCC channel.

### Construction of cell-cell communication subnetworks connected by ligand-receptor signaling

Discovering a pair of highly correlated GEMs across cell types suggests possible CCC between the cells expressing the GEMs. Subsequently, we conducted a systematic search and examination of potential CCC channels and networks. Using CRC pseudo-bulk RNA-seq data, we first searched for highly correlated GEMs (*GEM X* and *GEM Y*) that are expressed in different cell types (or clusters) to discover potential CCCs. In addition, we performed a causal analysis to find potential ligands and receptors that mediate CCCs. To that end, we then search for *LR* pairs such that *L* is co-expressed with *GEM X* and *R* is co-expressed with *GEM Y.* Finally, we assess whether conditioning on *LR*, the two marginally correlated GEMs become independent, which is denoted as *GEM X* 11 *GEM Y* | *LR*. Conditional independence is an important concept in causal analysis, and in our setting, if a pair of highly correlated GEMs becomes independent when conditioning on a LR pair, this LR pair is potentially involved in transmitting signals between cells expressing *GEM X* and *GEM Y*. Finding a *GEM X*-*LR*-*GEM Y* triplet leads to the hypothesis depicted in **Fig. 6A**. More specifically, a change in the state of the signaling pathway (*X*) in one cell promotes the release of the ligand and, at the same time, leads to transcriptomic changes represented by *GEM X*, which serves as a proxy of the pathway (*X*) state. The binding of the ligand with its receptor in another cell results in a cellular effect represented by the signaling pathway (*Y),* which leads to the changed gene transcription represented by *GEM Y*. Further extending such CCC analysis to multiple GEMs and cell types leads to a CCC network (CCCN), which may be used to understand how different cells reach homeostatic states in a TME through CCCs. If there are GEMs/LR pairs of interest, a CCC sub-network can be constructed based on the *GEM X*-*LR*-*GEM Y* triplets. We identified and illustrated a CCC sub-network related to immune checkpoint signaling in **Fig. 6B**. The network consists of T cells expressing TNK GEMs (TNK26, TNK35 and TNK50) and non-TNK cells expressing GEMs (Stroma37, B44, Mye21, Mye43, and Mye46), likely transmitted through PD-1, CTLA4 and CXCL13 signaling. In this network, the communication between cells expressing Mye9 (a combination of dendritic and macrophage cells) and TNK35 (LAYN^+^ Effector Treg) can be mediated through *CD274-PDCD1*; the communication between the cells expressing Mye43 (TAM: tumor-associated macrophage) and TNK35/TNK26 (PDCD1^+^CXCL13^+^ CD4^+^ and CD8^+^ T cells) is transmitted through *CD80/86-CTLA4*; the communication between Mye26-expressing NLRP3^+^INHBA^+^ macrophage cells and TNK50-expressing CXCL13 producing CD4^+^ T cells is facilitated via *CD274-PDCD1*; the communication between TNK50 and B44-expressing follicular B helper cell is transmitted through *CXCL13-CXCR5* which is consistent with the current literature ^44^. Moreover, we also discovered the communication between a specific subset of mCAF and effector Treg. The communication between Stroma37 expressing MMP1^+^CXCL5^+^mCAF cells and TNK35 expressing LAYN1^+^Treg cells is transmitted through *CD274-PDCD1*. *MMP1* ranks the top 3^rd^ gene associated with Stroma37 and has been shown to play an important role in CAF-induced protection and lead to therapy resistance ^45^. The communication between stromal cells and Tregs has been validated in several studies ^46^. Besides, TNK51 expressing Th1 cells communicate with myeloid GEMs (Mye8 and Mye42) expressing suppressor-like macrophage (M2) through *TNF*- *TNFRSF1B* ^47,48^. The above results demonstrate that re-constructing CCC networks provides insights into potential molecular mechanisms of CCCN underlying immune evasion mechanisms of tumors.

**Fig. 6:**
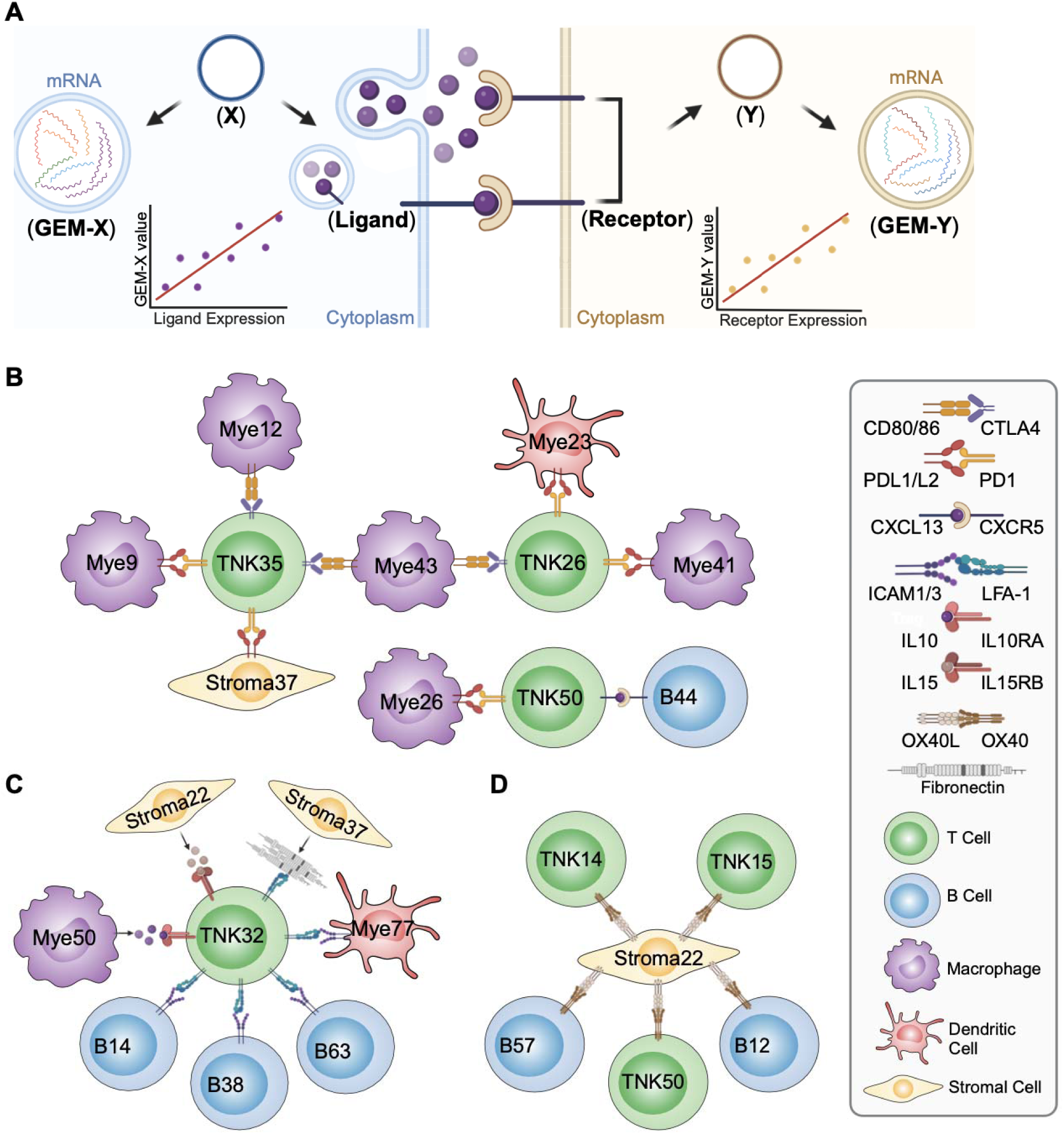
The newly discovered cellular networks associated with ligand-receptor interactions involved in the immune checkpoint inhibitor therapy. **A.** The framework of the conditional independence analysis to find the causal relationship between GEM-expressing cells transmitted by ligand-receptor signals. **B.** Subnetwork of intercellular communication related to PDL1/2-PD-1, CD80/CD86-CTLA4, and CXCL13-CXCR5 based on the conditional independence analysis. The detailed annotation of each GEM is listed in the interactive website. **C.** Subnetwork of intercellular communication related to TNK32 which represents the cellular function of NK cells. **D.** Subnetwork of intercellular communication between Stroma22 and GEMs from other cell types through the OX40L-OX40 signaling.

NK cells are a promising alternative platform for cellular immunotherapy, and clinical trials of NK cell immunotherapy have produced many encouraging clinical results ^49^. In **Fig. 6C**, a CCC sub-network illustrates the primary communications between TNK32-expressing NK cells and specific cellular functions of stromal, myeloid cells, and B cells through *ICAM1/3-LFA-1*, *IL10- IL10RA*, and *IL15-IL15RB* interactions. Besides, we discovered a highly ranked potential CCC channel involving Stroma22 expressing CAF, TNK14 expressing Treg and TNK50 expressing CXCL13^+^CD4^+^ T cells through *OX40L-OX40,* which is a costimulatory receptor as shown in **Fig. 6D**. Recent studies have shown the benefit of combining OX40 agonist and PD-1 inhibitor to improve the immunotherapy outcome ^50–52^. In both the CAMILLA trial dataset, where combined therapy involved a VEGFR2 inhibitor and anti-PD-L1, and the dataset from melanoma patients treated solely with anti-PD-1 immunotherapy, Stroma22 exhibits a notable difference between the responder and non-responder groups. (**Extended Data Fig. 7A**). In addition to the aforementioned findings, we have identified additional *GEM X-LR-GEM Y* triplets as potential CCCNs in colorectal cancer (CRC). These discoveries offer valuable insights into exploring intercellular communication within the TME and understanding the key cellular transcriptomic processes associated with ligand-receptor signaling. The details are listed in **Extended Data Table. 3**. To enhance the user-friendliness of our framework, we’ve created an interactive website where users can explore the annotation of each GEM, identify the top-ranked GEMs associated with each ligand-receptor pair, and investigate the *GEM X-LR-GEM Y* triplets of interest (http://44.192.10.166:3838/).

## Discussion

In this study, we showed that by deconvoluting the hierarchically organized transcriptomic programs and closely representing cell differential programs, the nHDP model can identify GEMs that are potentially regulated through CCC, discover possible CCC channels, and generate the hypothesis of the molecular mechanisms of CCC underlying heterogeneous tumor microenvironments of CRC. Detecting and representing the states of cellular signaling systems in individual cells is a fundamental task in single-cell biology. While there are different approaches for deconvoluting single-cell transcriptomic components ^18,53^, our results indicate that by modeling the hierarchical and compositional relationships among transcriptomic programs of cells, the nHDP model captures fine-grained transcriptomic signals better than other non-hierarchical models.

Although single-cell RNA-seq assays offer unparalleled cell-type-specific insights, their practical use is impeded by several obstacles (e.g. high cost of library preparation and sequencing) that hinder their broad application in clinical contexts. The evolution of bulk RNA-seq data over the past decades has yielded a substantial volume of samples conducive to the execution of pattern and machine learning investigations. The study of CCC, especially the TME associated with drug treatments, could be further promoted by integrating insights leaned from scRNA-seq data.

Therefore, effective methods for bulk RNA-seq deconvolution are still urgently needed. The GEMs learned from nHDP could be used as novel signatures for deconvoluting the specific transcriptomic processes in both bulk RNA-seq data and spatial transcriptomic data. Compared with two state-of-the-art statistical models, non-negative matrix factorization (NMF) and latent Dirichlet allocation (LDA), for identifying transcriptomic programs, nHDP model has the advantage of capturing the hierarchical structure of signaling pathways. GEMs derived from the nHDP model reflect the cellular states across multiple cell subtypes or specific to a particular cell subtype. Our recent study ^54^ showed that GEMs offered a more precise estimation of biological signals, such as immune cell infiltration, in comparison to the state-of-the-art deconvolution methods like Cibersort and SCNA ^55,56^. This improved accuracy enables the identification of different lymphocyte infiltration levels.

Identifying highly specialized GEMs enabled us to use the expression level of GEMs as proxies to represent the state of signaling pathways (unobserved) that regulate their expression. This, in turn, enabled us to search ligand-receptors that likely transmit extracellular signals to the pathways inside single cells, generating the hypothesis of how across-cell signaling influences intracellular signaling through CCC. Our results indicate that the *CD274-PDCD1* ligand-receptor pair may influence the state of the cellular signaling pathways that regulate the expression of certain members of TNK26 in exhausted T cells. The hypothesis is supported by multiple lines of evidence: The non-linear dose-response relationship between the approximated ligand-receptor complex and the GEM provides support from a biochemical point of view. The significant difference in TNK26 expression between responding and non-responding tumors treated with anti-PD-1 agents (both newly developed clinical trials in CRC and melanoma datasets) indicates that tumors with high TNK26 expression display active PD-L1-PD-1 signaling, rendering them more susceptible to the treatment. Conversely, tumors with low TNK26 expression exhibit relatively inactive PD-L1-PD-1 signaling, leading to their reduced responsiveness to the treatment. Based on these results, we hypothesize that the PD-L1-PD-1 signaling likely regulates some members of the TNK26. Further investigation of which members are regulated by this signal, e.g., using principled causal discovery methods, may enable researchers to delineate mechanisms of PD-1 signaling. Similar studies of the impact of other commonly co-expressed immune checkpoint proteins (receptors) in tumor-experienced T cells may reveal how different checkpoint signaling pathways cooperate and lead to immune suppression.

The discovery of a strong correlation of GEMs across cell types indicates that the expression of GEMs by different cell types is coordinated, and such coordination is likely due to CCC. We identified coordinated GEM expression patterns, referred to as CCC programs, and their spatial colocalization provides additional support for intercellular communication networks. We further noticed the differences in the activities of CCC programs may underlie the heterogeneity of TMEs, leading to tumor clusters that recapitulate well-appreciated molecular subtyping of CRC. Further investigation identifies ligand-receptors that likely transmit signals across GEMs in CCC channels.

We anticipate further investigating CCC channels active in each tumor/subcluster may provide mechanistic insights into the TME heterogeneity. In our population-wide exploration of CCC, we observed instances where multiple ligands could communicate with a single receptor, as illustrated by the TNK32 in **Fig. 6C**. In the causal theory, those ligands are multiple parents of the receptor. If we examine whether instantiated values of variables in a local network lead to context-specific independence, only certain parents may contribute to the state of children from the aspect of an individual tumor sample. For instance, in an individual tumor sample, only one type of cell among stromal, myeloid, or B cells may be involved in the interaction with TNK32- expressing cells. Our forthcoming efforts will involve employing the individualized Causal Bayesian Network (iCBN) model ^57^ to reveal the individualized CCCN within each tumor sample. This effort will contribute to a deeper understanding of the mechanisms underlying heterogeneity. Besides, this study approximates the ligand receptor interaction based on the product of [L] and [R] due to the feasibility. We realized that this approximation method using the gene expression is insufficient. However, since gene expression is one type of data available as a surrogate for ligand/receptor proteins, the method applied in this study is among the widely used approximation approach by the research community ^5,6^.

We envision that the framework presented in this study for deconvoluting single-cell transcriptomic programs to model CCC can be applied to various cancer types. Enhancing the nHDP model involves refining the initialization process and filtering variable genes, leading to a more effective capture of relationships within transcriptomic programs. This improvement will boost the model’s ability to detect GEMs regulated by specific transcriptomic programs. Future methodological developments for detecting fine-grained transcriptomic programs may involve leveraging recently developed topic modeling models to capture the compositionally co-expressed genes. Additionally, integrating single-cell RNA-seq with single-cell chromatin accessibility sequencing (scATAC) could aid in discovering active motifs associated with CCC networks. In conclusion, this study advances the understanding of CCCs mechanisms underlying heterogeneous TME and immune evasion mechanisms in CRC. These insights may uncover novel targets for immunotherapy and guide precision immunotherapy ^23^.

## Online Method

### Data Collection and Preprocessing

#### Single-cell transcriptomics

We collected single-cell RNA-seq data from 8 publicly available colorectal cancer data sources (**Extended Data Table. 1**) ^24,35,58,59^. The total number of patients is 153, of which there are 179 tumor samples and 100 normal samples. We retained cells that met the following criteria: nUMI > 500, nGene > 200, log10GenesPerUMI > 0.8, and mitoRatio < 0.2. To perform gene-level filtering, we retained only genes expressed in at least 0.1% of the cells. After the quality control (QC) analysis, we applied the total-count correction (library-size correction) and normalized the data matrix to 10,000 reads per cell. The annotations of cell major type were obtained from original publications and used for further analysis. We then divided cells into five major categories: 246,560 T and natural killer (TNK) cells, 94,663 myeloid cells, 96,637 plasma and B cells, 69,130 ^6^stromal cells, and 119,870 epithelial cells. When performing clustering analysis within each cell category, the Harmony algorithm ^60^ was applied to remove the batch effect, cell clustering was carried out using the Leiden algorithm ^61^, and the UMAP (Uniform Manifold Approximation and Projection) algorithm was used to visualize clusters ^62^. Seurat pipeline was used to find subclusters and their top variable genes within TNK, myeloid, plasmaB, stromal, and epithelial cells, respectively ^15,63^. The cell types were annotated with well-known marker genes: *CD3D, CD8A, FOXP3, IL7R, NKG7, GNLY* for T and NK cells; *CD79A, MS4A1, MZB1* for B and plasma cells; *CD14, CD68, CD1C, S100A8, C1QA, LYZ* for myeloid cells. The detailed markers we used were listed in **Extended Data Table. 5**.

### Bulk RNA-seq

The bulk RNA-seq cohorts collected in this study mainly serve for discovering potential GEM features strongly associated with the anti-PD-1 immunotherapy. Bulk RNA-seq data of 20 patients with metastatic CRC, treated with the combination of Cabozantinib plus durvalumab in the CAMILLA clinical trial and characterized by proficient microsatellite stable (MSS) tumors, were collected upon request ^28^. The number of responders and non-responders is 4 and 16, respectively. The responders include the CR (complete response) and PR (partial response) and the non-responders include the PD (progressive disease) and SD (stable disease). Three melanoma bulk RNA-seq data with response to anti-PD-1 treatment were downloaded from Liu et al, Riaz et al, and Gide et al for the validation analysis ^29–31^.

### Pseudo-bulk RNA-seq

To facilitate the clustering of GEMs and the searching for ligand-receptor pairs using the CRC single-cell data, we generated pseudo-bulk RNA-seq results for tumor samples (a *tumor-by-* matrix). For the tumor samples (n=151) that include both CD45^-^ and CD45^+^ cells, UMI counts of the same gene in each sample were aggregated, resulting in the expression data in raw count format. We then normalized the raw count expression data to log_2_(TPM+1).

### Spatial transcriptome

Spatial transcriptomic data of colorectal cancer and its corresponding liver metastasis using 10x Visium technology were collected from the study by Wu et al ^35^ and Valeolivas et al ^36^. These two cohorts contain 8 and 16 slides, respectively. As shown in **Extended Data Fig. 10**, these two cohorts contains 8 slides (Gao_colon1, Gao_colon2, Gao_colon3, Gao_colon4, Gao_liver1, Gao_liver2, Gao_liver3, and Gao_liver4) and 16 slides (SN048_A121573_Rep1, SN048_A121573_Rep2, SN048_A416371_Rep1, SN048_A416371_Rep2, SN84_A120838_Rep1, SN84_A120838_Rep2, SN84_A121573_Rep1, SN84_A121573_Rep2, SN123_A595688_Rep1, SN123_A798015_Rep1, SN123_A551763_Rep1, SN123_A938797_Rep1_X, SN124_A551763_Rep2, SN124_A938797_Rep2, SN124_A595688_Rep2, SN124_A798015_Rep2), respectively. We first used sctransform method ^64^ to normalize the raw count in each spot of the slide (a *spot-by-gene* matrix). The mean of top 50 genes associated with each GEM was then used to estimate the GEM values from normalized expression data. The deconvoluted spatial transcriptomic data is represented as a *spot-by-GEM* matrix. Function RCTD (R package spacexr) ^65^ was used to infer the cell-type composition in tumor region.

#### Identification of GEMs using nHDP

We used the nested hierarchical Dirichlet processes (nHDP) model ^54,66^ to identify patterns of GEMs from single-cell RNA-seq data. The flow of the training process is shown in **Fig. 1B & 1C**. We binarized the UMI count of each gene within a cell as the input of the nHDP model. If a gene is expressed in a cell with non-zero value, it will be assigned a binary value 1; Otherwise, it will be assigned a binary value of 0. A trained nHDP model returns two matrices: a cell-by-GEM count matrix, which collects the counts of UMIs assigned to a GEMs in each cell; a GEM-by-gene matrix, which represents the probability profile of genes in each GEM (**Fig. 1B**). We constructed a three-layer tree structure with the following branching factors: 5, 4, and 3 for 1^st^, 2^nd^, and 3^rd^ layers, respectively, as shown in **Fig. 1C**. In this model, the choice of tree structure (e.g., 5-4-3) was guided by biological priors such as the expected number of major lineages, ensuring that top level GEMs capture common programs while deeper nodes refine subtype- and function-specific signals. We systematically evaluated alternative tree structures (3-2-2, 4-3-2, 5-4-3) and demonstrated that larger and deeper trees yielded consistently lower perplexity and higher topic coherence, particularly as the training size increased^54^. The switching parameter γ was set to balance exploration of deeper nodes with probability ranging from 0 to 1, and mini-batch training with L1 K-means initialization further improved robustness against noise. To validate reproducibility, we performed simulation studies and tested independent datasets (PBMC 3K, 8K, 33K) repeatedly, finding consistent GEM structures and transferability across donors. Together, these analyses demonstrate that the nHDP model is robust to parameter choices when informed by biological context, and reproducible across independent runs and datasets.

The expression of GEMs (cell-by-GEM matrix) is added to the Seurat object as new features for plotting the UMAP of GEMs. The dominant function represented by a GEM is based on the rank of cell subtype percentage ("Distribution” in **Extended Data Fig. 4**). We used word clouds R package “wordcloud2” to visualize the probability profile of the top 20 associated genes of a GEM (**Fig. 1D**). We trained a nHDP model for each cell category (TNK, B, Mye, Stroma, and Epi) and retained GEMs expressed in over 1% of cells in a category for further analysis.

#### Functional analysis of GEMs

Transcription factor and enrichment analysis were applied to conduct functional analysis on GEMs. CHEA3 was used to conduct the transcription factor analysis ^25^. Reactome, KEGG and GO databases were used to conduct comprehensive pathway analysis using R package “msigdbr”^67^. Besides, we conducted a hypergeometric analysis ^68^ to annotate GEMs with well-known gene sets (H and C1-C8) listed in MsigDB ^69^. The significant enrichment was defined as p-value <=0.05 & FDR <= 0.05.

#### Estimation of GEM level in bulk and spatial RNA-seq data

To estimate the expression level of a GEM in bulk/pseudo-bulk/spatial RNA-seq data, we used two methods: 1) gene set variation analysis (GSVA) of the top 50 weighted genes in each GEM^70^. 2) mean expression of the top 50 weighted genes in each GEM.

### Fitting non-linear relationship between ligand-receptor pairs and GEMs

The ligand-receptor database was downloaded from Nichenet ^6,21,22^. To fit the dose-response curve between the expression of ligand-receptor and GEM value, we model the quantity of LR complex [LR] as the product of expression of L and R based on the fact that [LR] ∝ [L][R] using the pseudo-bulk RNA-seq data. We first scaled the GEM value to [0, 1] and then created a 4-parameter log-logistic function as 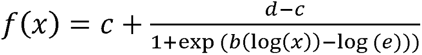, where *d* and *c* on 0 and infinite dose, *e* is the point of inflection and *b* being the hill slope using *drc/v3.0.1* ^71^. The coefficients and its associated *p*-values were estimated and adjusted with Benjamini-Hochberg procedure^22^. Sigmoid fitting with a significant adjusted *p*-value (<0.05) demonstrates a good fit between the expression of LR complex and the scaled GEM values.

### Assessing conditional independence of *GEM X*-*LR*-*GEM Y* triplets

To find out the ligand-receptor pairs that potentially mediate CCC between cells across cell categories expressing two correlated GEMs, conditional independence tests of *GEM X* GEM Y | LR were performed using the *bnlearn/v4.7.1* package in R ^72^. We estimated expression status of each GEM using the GSVA scores (*GSVA/v1.42.0*) ^70^ based on its top 50 genes. We calculated all pairwise correlations between GEMs expressed in different cell categories. If a pair of cross-cell-category GEMs exhibits significant correlation, we further search for LR pairs that rendered conditional independence (*GEM X* GEM Y | LR ). *P*-values of conditional independence tests were adjusted to control the false discovery rate using Benjamini-Hochberg method. We set the threshold of the conditional correlation between *GEM1* and *GEM2* at an adjusted *p*-value > 0.05 as conditional independent.

We further performed post-hoc proportional tests to remove false positive findings. For each conditional independent *GEM X—LR—GEM Y* triplet, we investigated the co-expression of *GEM X—L* in potential signal-originating cells and the co-expression of *R—GEM Y* in potential signal-receiving cells. To conduct the proportional tests, we first construct a 2 by 2 contingency table to measure the frequency of co-expression GEM and the tested gene. GEM and gene were considered presenting if the corresponding values are beyond 0 and not presenting otherwise. Then the statistics of proportional tests 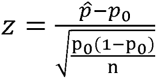 were estimated to test if the proportion of co-expressing cells is significantly higher than the proportion of not co-expressing cells, where p is the proportion of cells that co-express GEM and tested gene and *p_0_* is the proportion of cells that not co-express GEM and tested gene, *n* is the total number of cells that express such GEM. *p*-value of each post-hoc test was also adjusted using the Benjamini-Hochberg method for false discovery rate control ^73^. Ligand-receptor pairs that passed both conditional independence test and post-hoc proportional test were considered valid for downstream analysis.

To reduce false positives, we applied multiple criteria to filter GEM pairs. At the pseudo-bulk level, the correlations of *GEM X–GEM Y*, *GEM X–L*, and *R–GEM Y* were all required to exceed 0.3. In addition, we examined expression at the single-cell level to confirm the presence of specific molecules (ligand/receptor genes) in GEM-expressing cells, thereby further reducing false positives. Specifically, we required that more than 10% of cells expressing GEM X also express ligand gene *L*, and more than 10% of cells expressing *GEM Y* also express receptor gene R.

#### Assessing Spatial Colocalization

Spatial colocalization was evaluated by calculating spot-wise spatial correlations. To account for soluble ligands, we assigned a weight of 1 to the center spot and first-layer neighbors, and a weight of 0.5 to second-layer neighbors. Because each spatial spot contains a mixture of cells, we inferred the cell composition within each spot (with the percentages of different cell types summing to 1) using RCTD ^65^.

We then multiplied the GEM value by the cell type percentage to estimate the expression of GEMs for a specific cell type. We calculated five types of correlations: (1) *GEM X-L* (ligand), (2) *GEM X-GEM Y*, (3) *R* (receptor)-*GEM Y*, (4) *GEM X*-*LR*, and (5) *LR*-*GEM Y*. For *GEM X-L* and *GEM X-GEM Y* correlations, we used lagged *GEM X* values derived from the two layers of neighboring spots. For *GEM X-L* and *R-GEM Y* correlations, we first computed the ligand– receptor product using the lagged ligand value, and then computed spot-wise correlation between ligand–receptor product and *GEM X* or *GEM Y*. Statistical significance of each correlation was assessed using Pearson’s correlation test to obtain p-values. Spatial pairs were considered colocalized only if the correlations of *GEM X-GEM Y*, *GEM X-L* and *LR-GEM Y* all exceed 0.3.

We then multiplied the GEM value by the cell type percentage to estimate GEM expression for a specific cell type. We calculated five types of correlations: (1) *GEM X–L* (ligand), (2) *GEM X– GEM Y*, (3) *R* (receptor)–*GEM Y*, (4) *LR–GEM Y*. For *GEM X–L* and *GEM X–GEM Y* correlations, we used lagged GEM X values derived from two layers of neighboring spots. For *LR–GEM Y* correlations, we first computed the ligand–receptor product using the lagged ligand value, and then calculated spot-wise correlations between the ligand–receptor product and *GEM Y*. The statistical significance of each correlation was assessed using Pearson’s correlation test to obtain p-values. Spatial pairs were considered colocalized only if the correlations of *GEM X– GEM Y*, *GEM X–L*, and *LR–GEM Y* all exceeded 0.3.

#### Identifying programs of GEMs

Hierarchical clustering was performed on a Euclidean distance matrix ^74^ to identify GEM programs using the pseudo-bulk expression of GEMs (sample-by-GEM matrix) as input.

#### Evaluating GEMs as potential biomarkers for anti-PD-1 therapies

We performed GSVA analysis to deconvolute the expression levels of GEMs for each bulk tumor sample. Samples were divided into the response group if they showed complete response (CR) or partial response (PR) to anti-PD-1 whereas non-response group if progressive disease (PD) or stable disease (SD) were observed. We employed the one-sided Wilcoxon rank test ^75^ to identify GEMs that exhibited a significant difference in expression levels between these two groups.

## Availability

nHDP model designed for analyzing single-cell transcriptome (scGEM): https://github.com/hansolo-bioinfo/scGEM

CRC Atlas: https://github.com/hansolo-bioinfo/CRCAtlas

Interactive website: http://44.192.10.166:3838/

## Ethics

No animal or human subjects is involved in this study.

## Acknowledgments

This research was supported in part by The National Library of Medicine, National Human Genome Research Institute, and National Cancer Institute at the National Institutes of Health grants R00LM013089, R01HG014023, R01LM012011 and R01CA254274. The Department specifically disclaims responsibility for any analysis, interpretations or conclusions.

## Supplementary Figures

**Supp Fig. 1:**
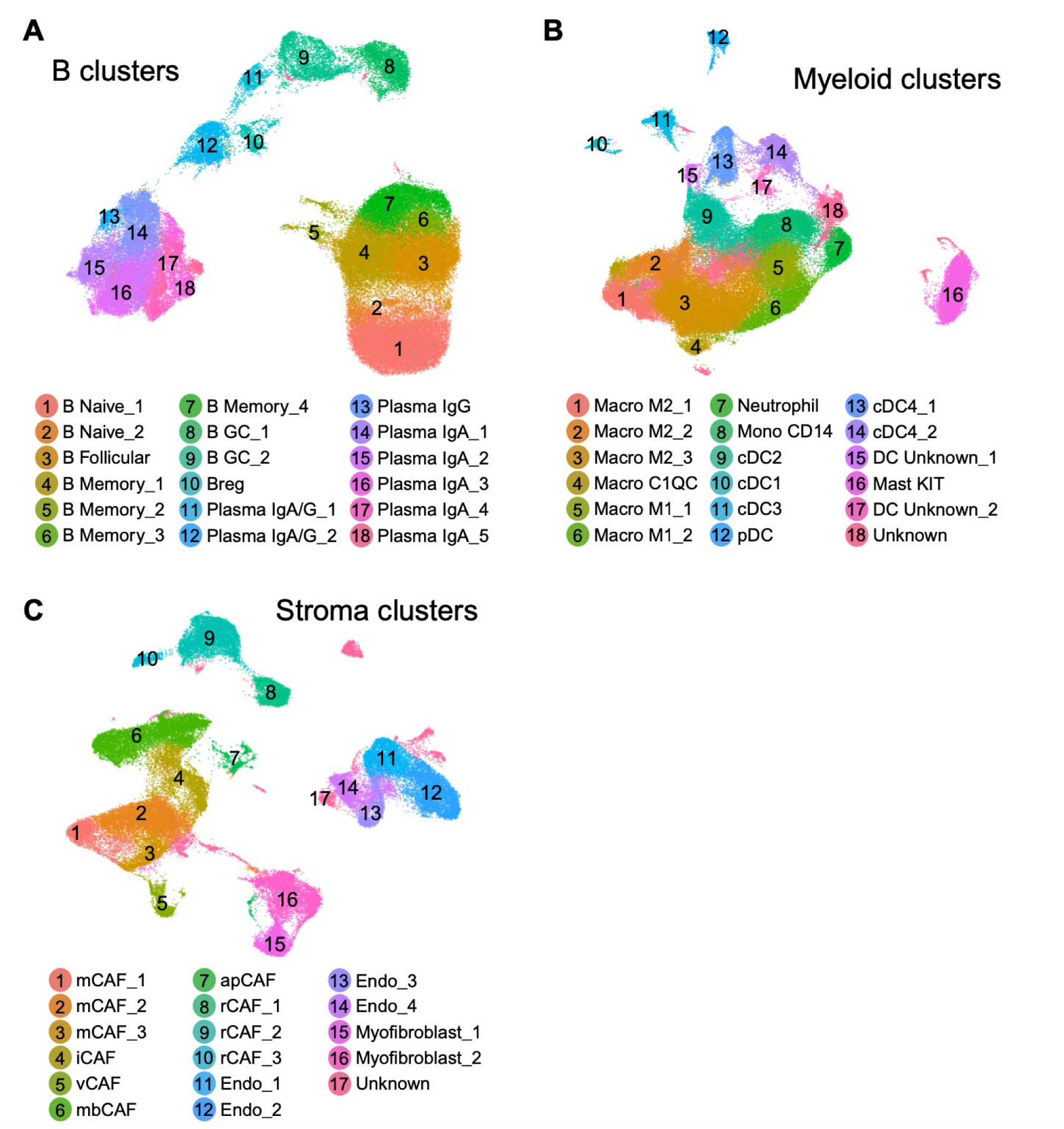
UMAP of B, Myeloid and Stromal cells. **A.** The cell clusters for B and Plasma cells. **B.** The cell clusters for myeloid cells. **C.** The cell clusters for stromal cells.

**Supp Fig. 2:**
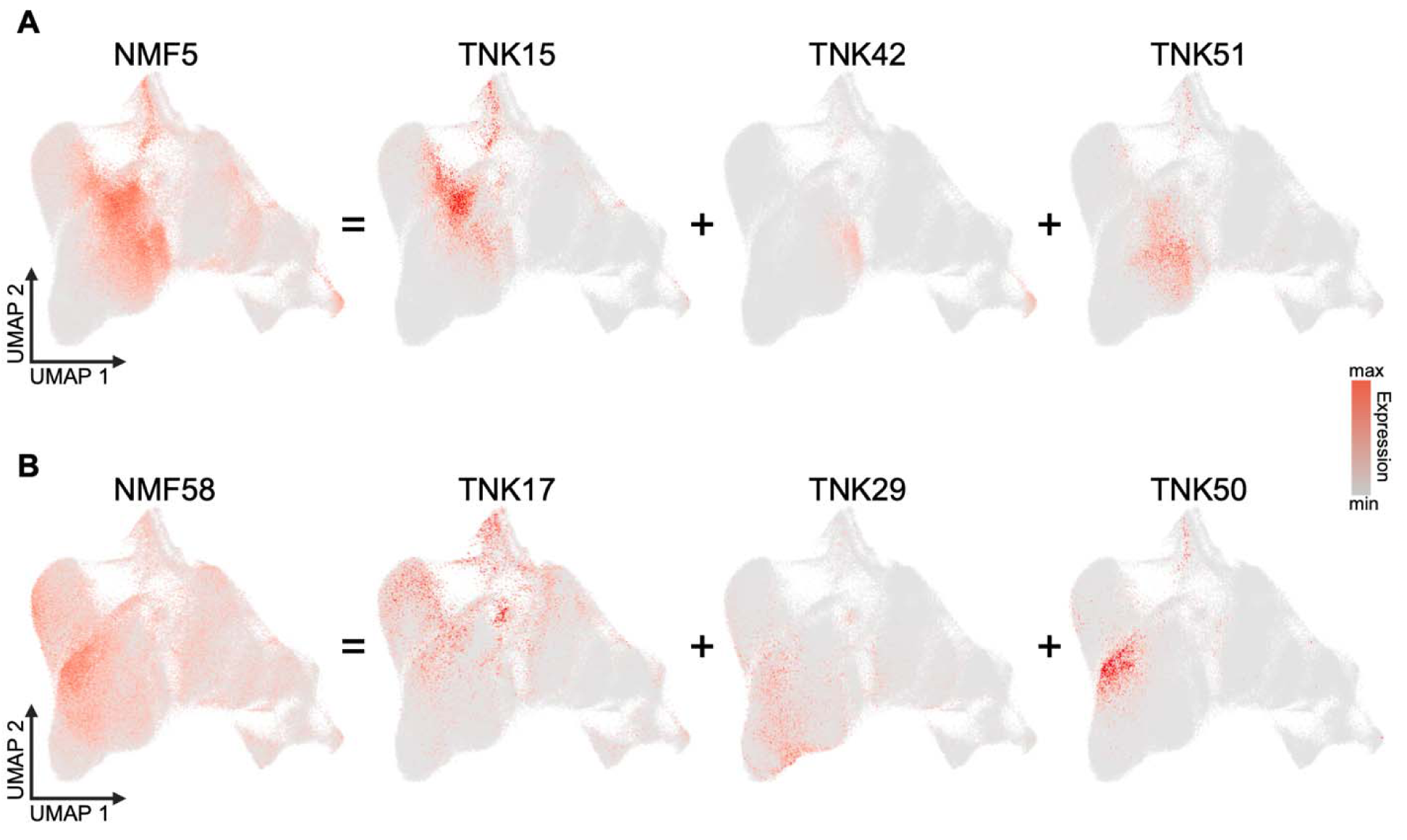
Comparison between GEMs learned from nHDP and GEMs learned from NMF. **A.** The cells expressing NMF5 are the combination of cells expressing TNK15, TNK42 and TNK51. **B.** The cells expressing NMF58 are the combination of cells expressing TNK17, TNK29, and TNK50.

**Supp Fig. 3:**
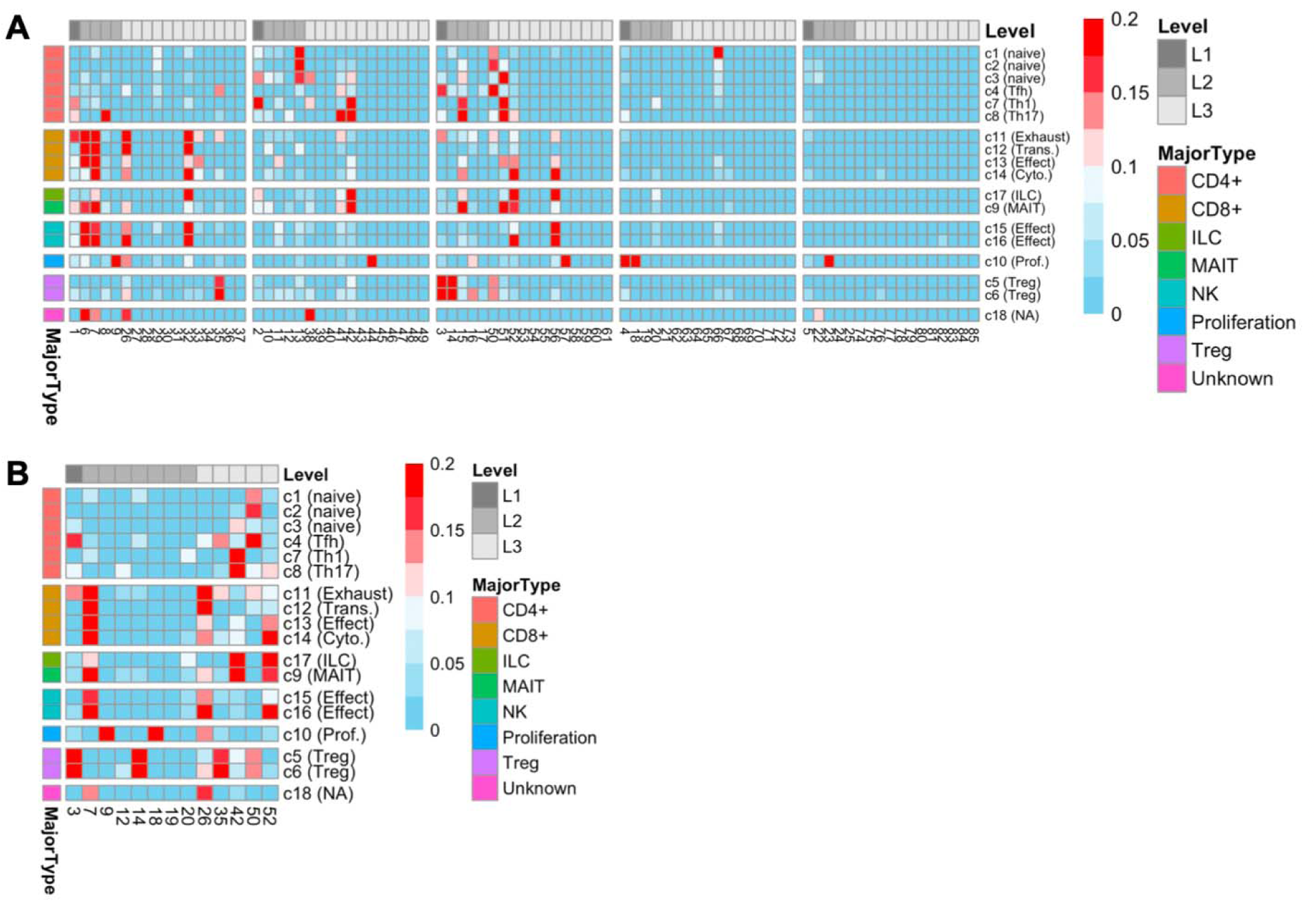
Cosine similarity between top50 GEM associated genes and highly expressed marker genes in T cell subtypes. **A.** Heatmap of cosine similarities between differentially expressed genes of T cell subtype and top 50 genes of all 85 T cell GEMs learned from nHDP model. GEMs are labeled with respect to their tree levels (L1, L2, and L3) and major T cell subtypes including CD4^+^, CD8^+^, ILC, MAIT, NK, Treg and proliferation T cells. **B.** The heatmap of cosine similarity for significant T cell GEMs summarized in Fig. 2E are shown.

**Supp Fig. 4:**
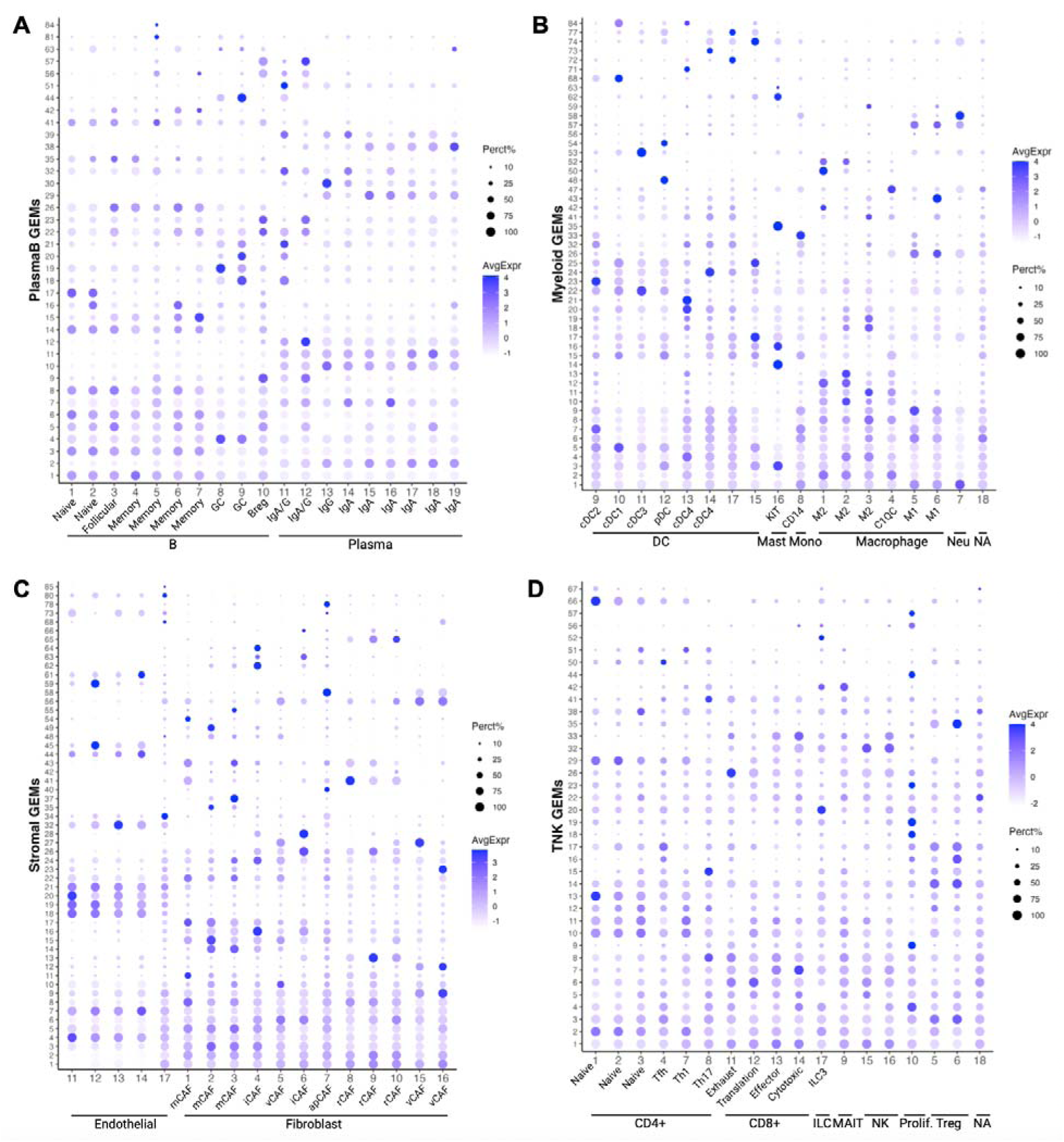
The dot plot comparing the similarity between GEMs and traditional cell clusters. **A.** The dot plot for B and Plasma cells. **B.** The dot plot for myeloid cells. **C.** The dot plot for stromal cells. **D.** The dot plot for TNK cells. The color and size of the dot reflect the average expression and percentage of overlapped cells, respectively.

**Supp Fig. 5:**
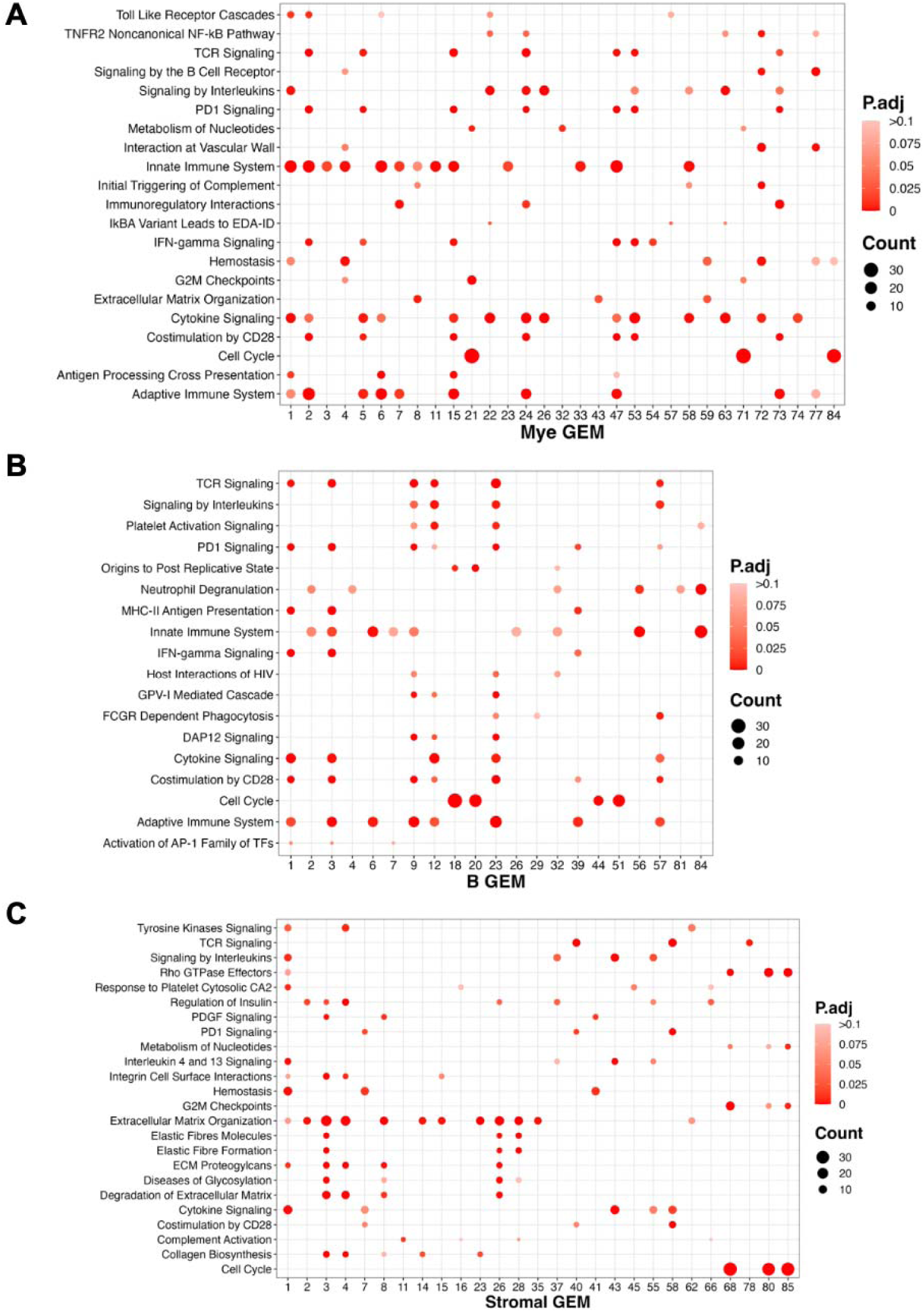
The enriched signaling pathways. **A.** The enriched signaling pathway for myeloid cells. **B.** The enriched signaling pathway for B and plasma cells. **C.** The enriched signaling pathway for stromal cells.

**Supp Fig. 6:**
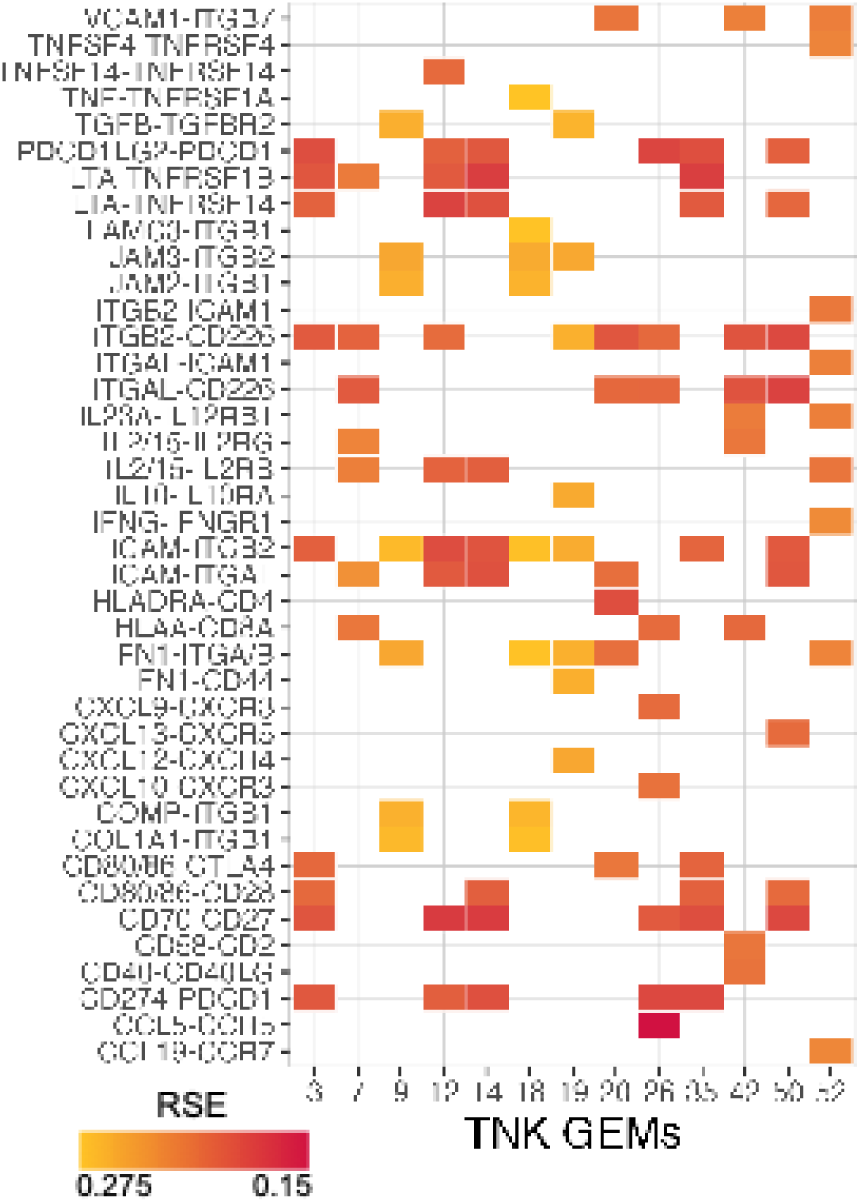
The summarization of top ligand-receptor pairs significantly associated with TNK GEMs. The residual standard error (RSE) of the sigmoid fitting model against ligand-receptor expression production and GEM signals are shown in gradient color. Ligand-receptor pairs with significant p-value of sigmoid model are displayed.

**Supp Fig. 7:**
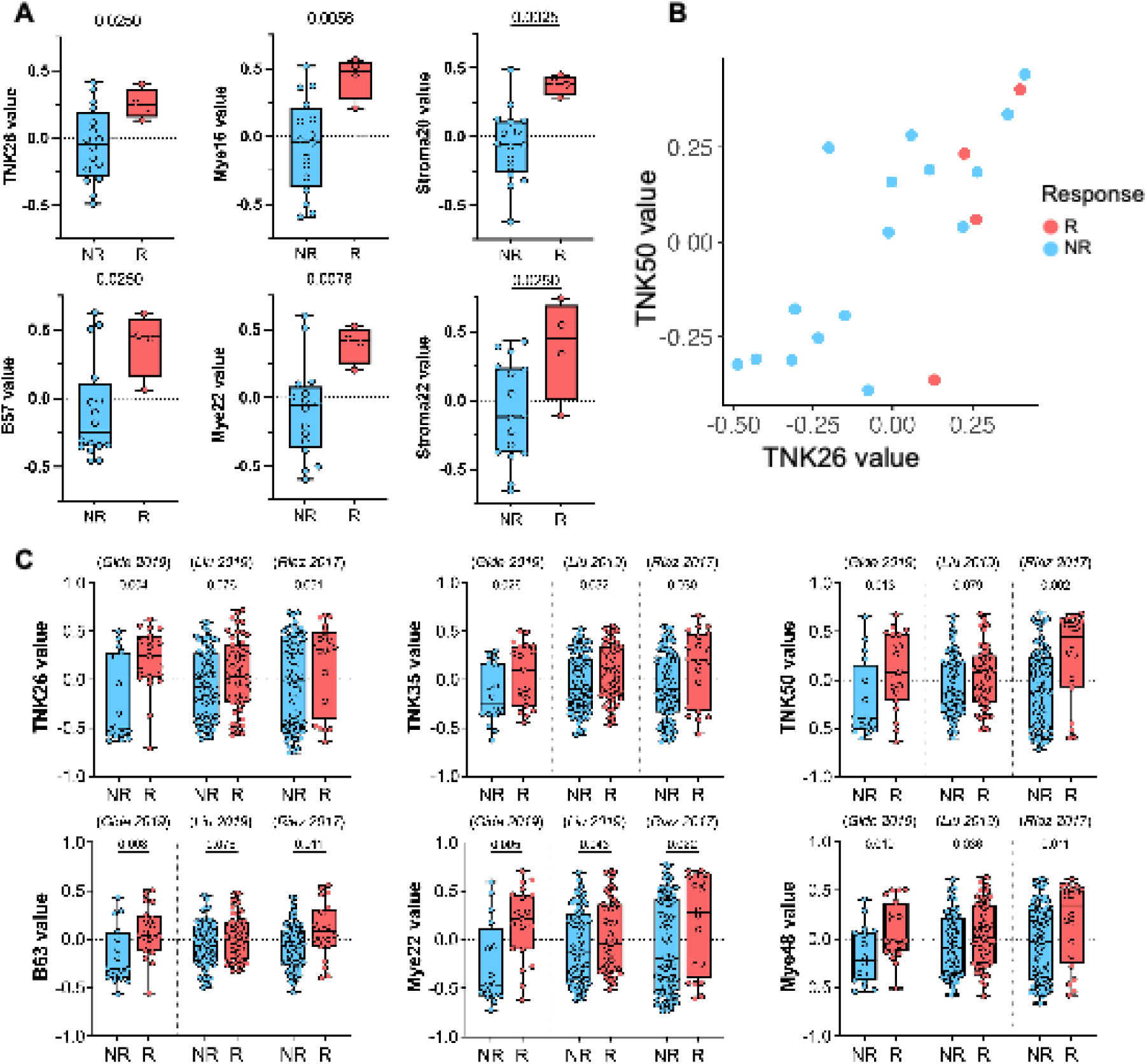
Example of GEMs significantly differentially expressed in responder group and non-responder group. **A.** Example of GEMs significantly differentially expressed in responder group and non-responder group treated with Cabozantinib plus durvalumab (CAMILLA trial). **B.** The correlation between TNK26 and TNK50 in CAMILLA cohort. **C.** Example of GEMs significantly differentially expressed in responder group and non-responder group treated with solo anti-PD-1 immunotherapy (three independent melanoma datasets).

**Supp Fig. 8:**
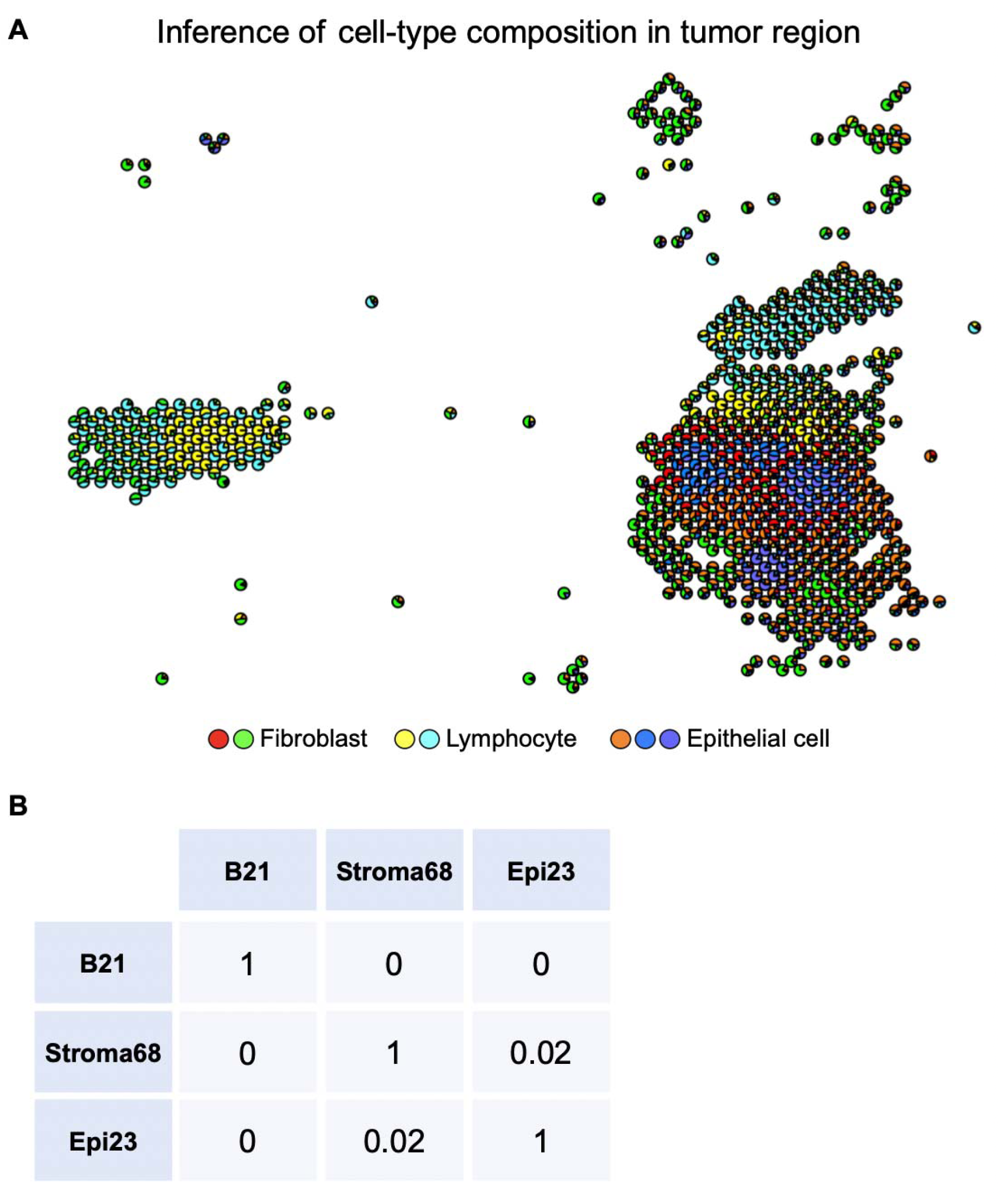
Exploring the uniqueness of three GEMs in CCC program 2. **A.** The inference of cell-type composition in tumor region of Fig. 5D. **B.** The cosine similarity between B21, Stroma68 and Epi23 using the expression level of the top 50 associated genes.

**Suppl Fig. 9:**
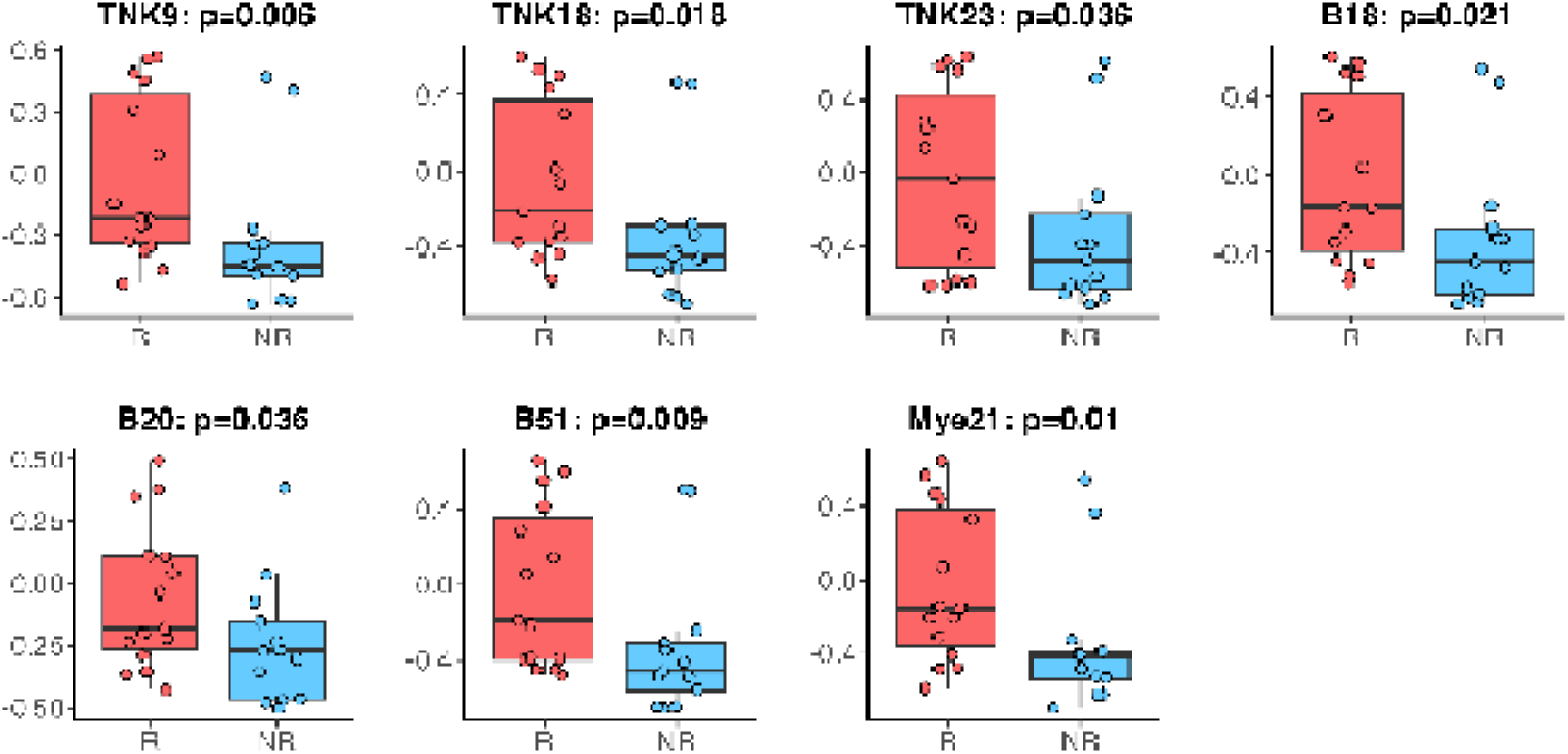
Example of GEMs in the proliferation program (CCC program2), which are significantly differentially expressed in responders and non-responders treated with regorafenib and nivolumab combination therapy.

**Suppl Fig. 10:**
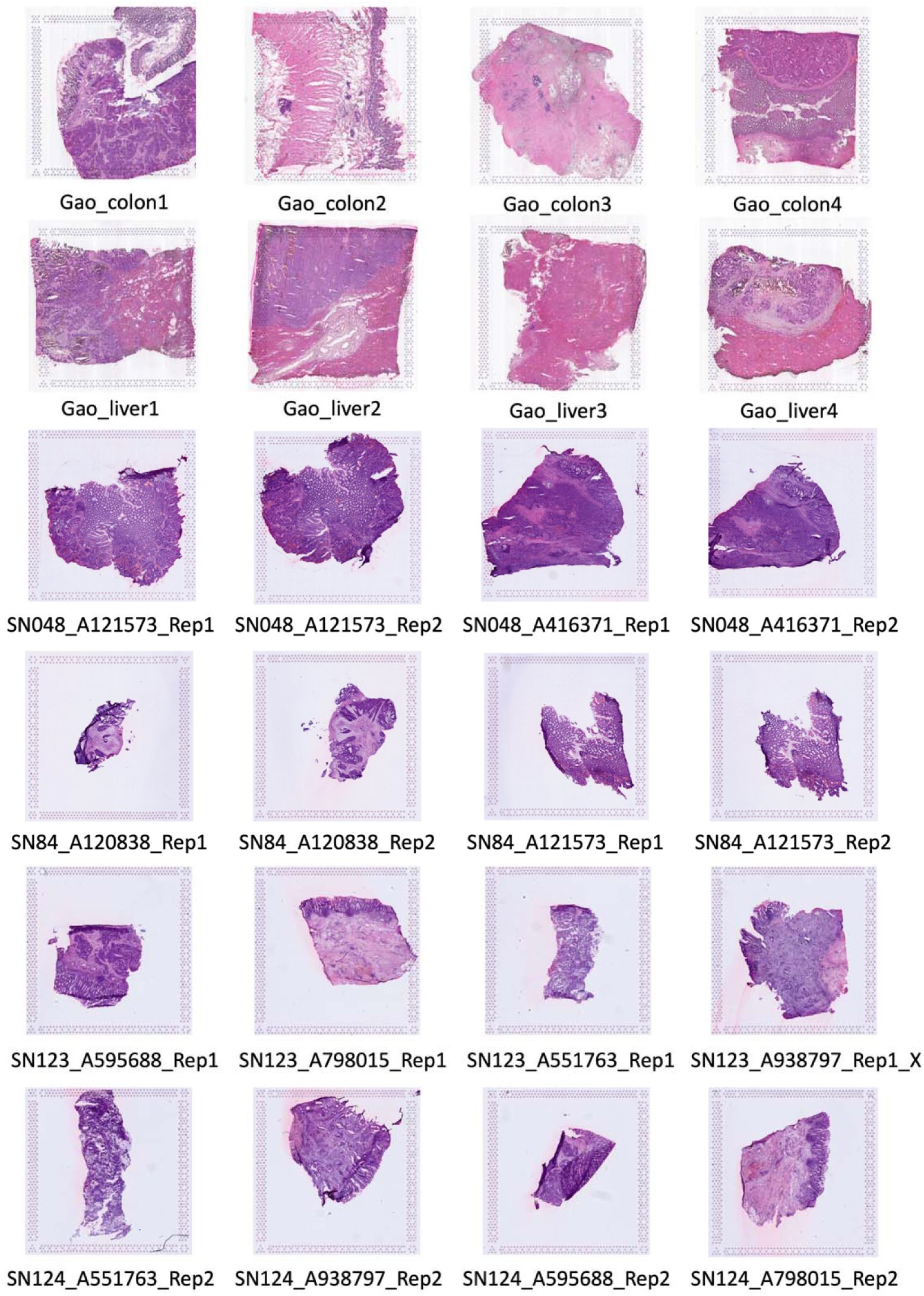
HE slides in the two publicly available spatial transcriptomics cohorts used in this study.

